# Nuclear RNA binding regulates TDP-43 nuclear localization and passive nuclear export

**DOI:** 10.1101/2021.08.24.457459

**Authors:** Lauren Duan, Benjamin L. Zaepfel, Vasilisa Aksenova, Mary Dasso, Jeffrey D. Rothstein, Petr Kalab, Lindsey R. Hayes

## Abstract

Nuclear clearance of the DNA/RNA-binding protein TDP-43 is a pathologic hallmark of amyotrophic lateral sclerosis and frontotemporal dementia that remains unexplained. Moreover, our current understanding of TDP-43 nucleocytoplasmic shuttling does not fully explain the predominantly nuclear localization of TDP-43 in healthy cells. Here, we used permeabilized and live-cell models to investigate TDP-43 nuclear export and the role of RNA in TDP-43 localization. We show that TDP-43 nuclear efflux occurs in low-ATP conditions and independent of active mRNA export, consistent with export by passive diffusion through nuclear pore channels. TDP-43 nuclear residence requires binding to GU-rich nuclear intronic pre-mRNAs, based on the induction of TDP-43 nuclear efflux by RNase and GU-rich oligomers and TDP-43 nuclear retention conferred by pre-mRNA splicing inhibitors. Mutation of TDP-43 RNA recognition motifs disrupts TDP-43 nuclear accumulation and abolishes transcriptional blockade-induced TDP-43 nuclear efflux, demonstrating strict dependence of TDP-43 nuclear localization on RNA binding. Thus, the nuclear abundance of GU-rich intronic pre-mRNAs, as dictated by the balance of transcription and pre-mRNA processing, regulates TDP-43 nuclear sequestration and availability for passive nuclear export.

## Introduction

Transactive response DNA binding protein 43 kD (TDP-43) is an essential DNA/RNA-binding protein that plays a major role in RNA processing and stability (reviewed in Prasad et al., 2019; François-Moutal et al, 2019). Loss of nuclear TDP-43 expression and accumulation of cytoplasmic aggregates is a pathologic hallmark of amyotrophic lateral sclerosis (ALS) and frontotemporal dementia (FTD) (Neumann *et al*, 2006; Arai *et al*, 2006), where approximately 97% and 45% of cases show TDP-43 pathology at autopsy, respectively (reviewed in Ling et al., 2013). Substantial evidence links TDP-43 disruption to the pathogenesis of ALS and FTD, via loss of nuclear splicing regulation (Polymenidou *et al*, 2011; Tollervey *et al*, 2011; Ling *et al*, 2015) and the toxic effects of cytoplasmic aggregates (reviewed in Vanden Broeck et al., 2014; Prasad et al, 2019). Mutations in TDP-43 have been identified in families with inherited ALS (Kabashi *et al*, 2008; Sreedharan *et al*, 2008; Deerlin *et al*, 2008; Gitcho *et al*, 2008) and FTD (Borroni *et al*, 2009; Kovacs *et al*, 2009) and cause neurodegeneration in disease models (reviewed in Buratti, 2015), supporting a role for TDP-43 disruption in the evolution of disease. Remarkably, the vast majority of ALS and FTD cases involving TDP-43 cytoplasmic mislocalization are not associated with TDP-43 mutations, indicating that diverse genetic, environmental, or age-related causes, possibly all involved in the same underlying process, drive the nuclear clearance of TDP-43 in disease. However, the factor(s) responsible for TDP-43 nuclear accumulation in healthy cells and mislocalization in ALS/FTD remain unclear.

TDP-43 is a member of the heterogenous ribonucleoprotein (hnRNP) family of RNA binding proteins (RBPs) that contains an N-terminal bipartite nuclear localization signal (NLS), two RNA recognition mofits (RRM1 and RRM2), and an intrinsically disordered C-terminal domain (reviewed in François-Moutal et al, 2019). RNA crosslinking and immunoprecipitation (CLIP-seq) studies show that TDP-43 preferentially binds GU-rich RNA motifs, particularly within introns (Polymenidou *et al*, 2011; Tollervey *et al*, 2011), consistent with its essential role in regulating alternative splicing and repression of cryptic exons (Ling *et al*, 2015).

Like many hnRNPs, TDP-43 continuously shuttles between the nucleus and cytoplasm (Ayala *et al*, 2008). Ran-regulated active TDP-43 nuclear import occurs via binding of its NLS to importins α and β (Ayala *et al*, 2008; Nishimura *et al*, 2010). A modest degree of passive nuclear import also likely occurs based on size and the low level of persistent nuclear entry seen in cells expressing TDP-43-ΔNLS (Ayala *et al*, 2008; Winton *et al*, 2008). TDP-43 nuclear export was initially thought to occur via the exportin-1 (XPO1) receptor and a putative nuclear export signal (NES) in RRM2 (Winton *et al*, 2008). However, several groups recently showed that NES deletions do not disrupt TDP-43 export, nor does XPO1 knockdown or inhibition by selective inhibitors of nuclear export (Ederle *et al*, 2018; Pinarbasi *et al*, 2018; Archbold *et al*, 2018).

Adding 54-119 kD tags to TDP-43 strongly slowed its export, as would be expected for a passive export mechanism rather than an energy-dependent, active process (Pinarbasi *et al*, 2018; Ederle *et al*, 2018). Meanwhile, transcriptional blockade promotes TDP-43 nuclear efflux (Ayala *et al*, 2008; Ederle *et al*, 2018), as does overexpression of the mRNA export receptor NXF1 (Archbold *et al*, 2018), suggesting a role for nuclear RNA in mediating TDP-43 nuclear localization and raising the possibility of TDP-43 co-export with mRNA, perhaps via the TREX (TRanscription/EXport) pathway (discussed in Ederle & Dormann, 2017).

Here, we aimed to elucidate the mechanism of TDP-43 nuclear export and test the hypothesis that TDP-43 nuclear residence depends on its preferential binding to nuclear RNAs. Consistent with this idea, we found that RNase-mediated nuclear RNA degradation in permeabilized cells markedly disrupted TDP-43 nuclear sequestration, allowing TDP-43 to diffuse from the nucleus in low-ATP conditions. Moreover, in permeabilized and live cell assays, introduction of GU-rich oligomers induced TDP-43 exit from the nucleus, likely by competitive dissociation from endogenous nuclear RNAs. Inhibition of pre-mRNA splicing caused dose-dependent TDP-43 nuclear accumulation and prevented its nuclear export upon transcriptional blockade, further supporting the notion that nuclear accumulation of TDP-43 depends on its association with pre-mRNAs. Taken together, our results indicate that binding to GU-rich intronic pre-mRNAs retains TDP-43 within nuclei and dictates its availability for passive diffusion from the nucleus.

## Results

### NVP-2-induced RNA Pol II inhibition promotes rapid TDP-43 nuclear export

Actinomycin D, a DNA-intercalating agent and pan-transcriptional inhibitor, has been shown to induce the nuclear efflux of a subset of nuclear RNA-binding proteins (RBPs) including hnRNPA1 (Piñol-Roma & Dreyfuss, 1992), SR proteins (Cáceres *et al*, 1998), and TDP-43 (Ayala *et al*, 2008; Ederle *et al*, 2018). To confirm that TDP-43 nuclear accumulation is sensitive to inhibition of mRNA synthesis and avoid the possible pleiotropic effects of DNA intercalation, we treated HeLa cells with NVP-2, an inhibitor of CDK9-dependent RNA Polymerase II activation (Olson *et al*, 2018) and monitored TDP-43 localization by immunostaining (**Fig 1A-C**). RNA synthesis was analyzed in parallel via a brief pulse of 5-ethynyl-uridine (5-EU) prior to fixation, to facilitate ‘click-chemistry’ labeling of newly transcribed RNAs (Jao & Salic, 2008). Cells were imaged with an automated high-content spinning disc confocal microscope, and a translocation algorithm was used to quantify the background-corrected fluorescence intensity in the nuclear and cytoplasmic compartments (Hayes *et al*, 2020). Like actinomycin D (**Suppl fig 1A-C**), NVP-2 caused progressive, dose-dependent inhibition of RNA synthesis (**Fig 1A-B**), sparing RNA puncta within nucleoli resulting from RNA polymerase I-dependent ribosomal RNA synthesis (**Fig 1A** arrows) (Sharifi & Bierhoff, 2018). In parallel, there was progressive TDP-43 nuclear efflux, as measured by a decrease in the nuclear to cytoplasmic (N/C) ratio (**Fig 1A,C**). Importantly, the drop in the TDP-43 N/C ratio induced by both NVP-2 and actinomycin D corresponded to a decrease in nuclear immunofluorescence and an increase in cytoplasmic immunofluorescence (**Suppl fig 1D-E**), consistent with TDP-43 translocation from the nucleus to the cytoplasm. NVP-2 and actinomycin D treatment of mouse primary cortical neurons also induced time- and dose-dependent transcriptional inhibition and TDP-43 nuclear efflux (**Suppl fig 1F-J**).

**Figure 1.**
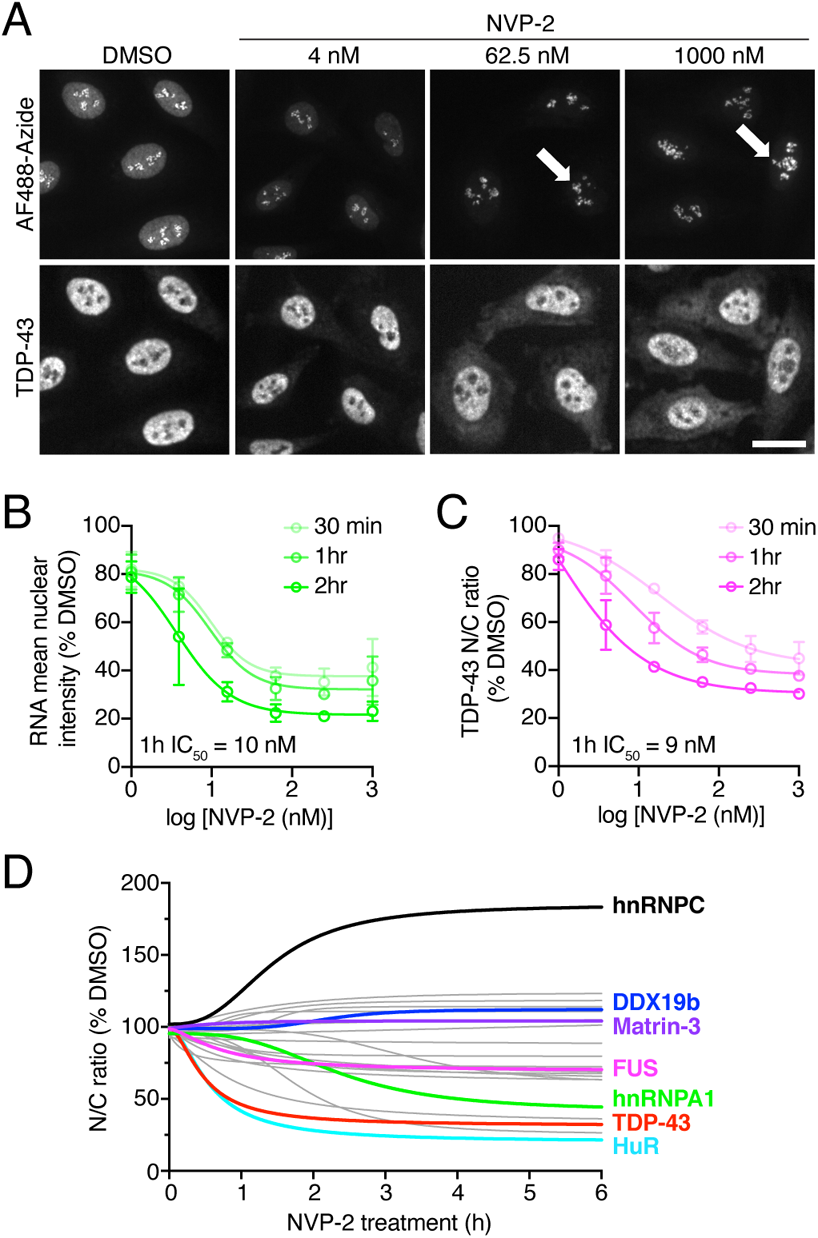
NVP-2-induced RNA Pol II inhibition promotes TDP-43 nuclear export. A. Newly-synthesized RNAs labeled with AF488-picolyl azide via 5-EU incorporation/‘click chemistry’ (top) and TDP-43 immunofluorescence (bottom) in HeLa cells treated with DMSO (vehicle) vs. NVP-2 for 1 h. Arrows indicate rRNA puncta unaffected by NVP-2. Scale bar = 25 µm. B, C. High content analysis of nascent RNA (AF488-azide) mean nuclear intensity (B) and TDP-43 nuclear to cytoplasmic (N/C) ratio (C) expressed as percent DMSO control. N = mean of 2442 cells/well in each of 3 independent biological replicates. Mean ± SD is shown. The IC_50_ for 1 h treatment is indicated, as calculated by non-linear regression. D. RBP N/C ratios in HeLa cells treated with 250 nM NVP-2 for 30 min to 6 h. Selected proteins are labeled (the full panel is detailed in **Suppl fig 2**). N = mean of 2716 cells/well. Curves were fit by non-linear regression using the mean of 2-4 biological replicates per RBP.

Next, we compared the effect of NVP-2 on the localization of TDP-43 versus a panel of other RBPs, including hnRNPs, RNA export proteins, members of the exon-junction complex, and splicing-associated proteins (**Fig 1D and Suppl fig 2A-B**). Cells were treated with NVP-2 for up to 6 h, fixed, and RBP localization was analyzed by immunofluorescence and automated high-content analysis. RBP responses were diverse, including rapid nuclear efflux (TDP-43, HuR), slow nuclear efflux (FUS, hnRNPA1), and no change (Matrin-3). DDX19b, an ATP-dependent RNA helicase in the mRNA export pathway that is tethered to the NPC (Napetschnig *et al*, 2009; Hodge *et al*, 2011) was unchanged, serving as an internal control for nuclear stability. hnRNPC, a resident nuclear hnRNP that does not exhibit N/C shuttling (Piñol-Roma & Dreyfuss, 1992; Ederle *et al*, 2018) showed nuclear accumulation. Interestingly, TDP-43 was among the most highly responsive RBPs to transcriptional blockade, suggesting strict dependence on nascent nuclear RNA levels.

**Figure 2.**
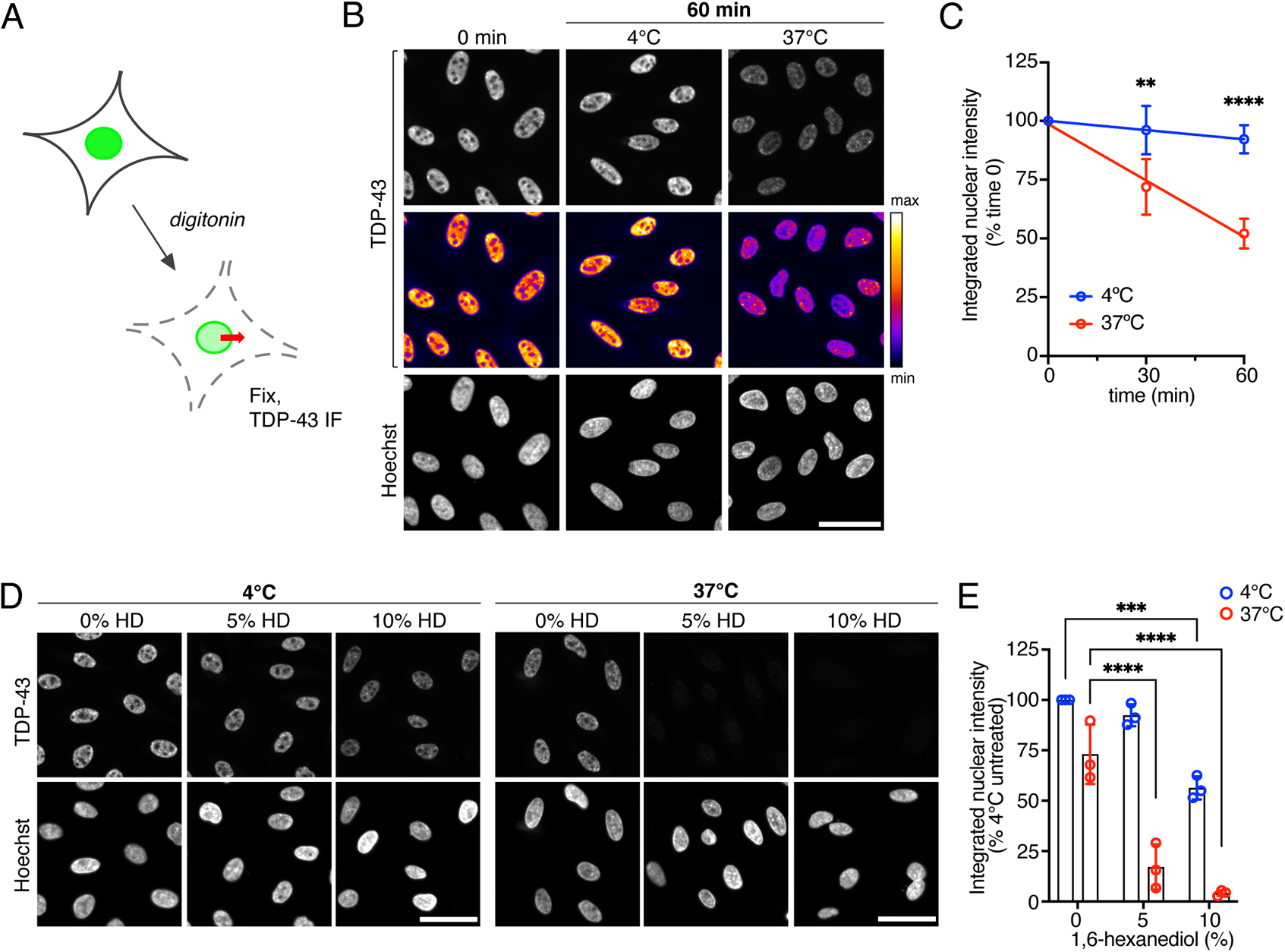
TDP-43 exits the nucleus by passive diffusion in permeabilized cells. A. Schematic of permeabilized cell TDP-43 passive nuclear export assay. IF = immunofluorescence. B. TDP-43 immunofluorescence (top) and Hoechst nuclear stain (bottom) in permeabilized HeLa cells fixed immediately post-permeabilization (0 min) and 60 min post-permeabilization, following incubation at 4°C or 37°C. The intensity histogram for all TDP-43 images was normalized relative to the 0 min control, prior to application of the pseudo-color LUT (middle) to illustrate relative TDP-43 nuclear intensity. Scale bar = 50 μm. C. Integrated nuclear intensity of TDP-43 in permeabilized HeLa cells over time at 4°C vs. 37°C expressed as percent of time 0. N = mean of 2185 cells/well/condition in each of 4 independent biological replicates. D. TDP-43 immunofluorescence staining (top) and Hoechst nuclear stain (bottom) in permeabilized HeLa cells incubated for 30 minutes post-permeabilization at 4°C vs. 37°C with increasing concentrations of 1,6-hexanediol. The intensity histogram for all TDP-43 images was normalized relative to 0% HD (4°C). Scale bar = 50μm. E. Integrated nuclear intensity of TDP-43 in permeabilized HeLa cells incubated for 30 minutes at 4°C vs. 37°C with increasing concentrations of 1,6-hexanediol. Data are expressed as percent untreated, 4°C. N = mean of 2389 cells/well/condition in each of 3 independent biological replicates. In C, E, **p<0.01, ***p<0.001, ****p<0.0001 by 2-way ANOVA with Tukey’s multiple comparisons test.

### TDP-43 exits the nucleus by passive diffusion

Recent studies showed that TDP-43 nuclear export is independent of the XPO1 nuclear export pathway (Archbold *et al*, 2018; Pinarbasi *et al*, 2018; Ederle *et al*, 2018) and may passively leave the nucleus in a size-limited manner (Pinarbasi *et al*, 2018; Ederle *et al*, 2018). In support of this, we found that the addition of a relatively small 27-kD YFP tag markedly inhibited NVP-2-induced TDP-43 nuclear efflux (**Suppl fig 1K**). However, the mechanism of passive versus active TDP-43 export across the NPC remains to be confirmed. Permeabilized cell assays have been widely used to study nucleocytoplasmic transport, facilitating investigations of transport mechanisms in isolation from other cellular processes and permitting a broad range of experimental perturbations (Adam *et al*, 1990; Cassany & Gerace, 2009). Most commonly, digitonin is used to selectively perforate the plasma membrane (Colbeau *et al*, 1971; Adam *et al*, 1990), releasing the cytoplasm and leaving the nuclear membrane and NPCs intact and able to perform either passive or energy-dependent bidirectional transport (Hayes *et al*, 2020). Here, we developed an assay for the passive nuclear export of endogenous TDP-43 in digitonin-permeabilized HeLa cells (**Fig 2A**). Post-permeabilization, cells were incubated in energy-free buffer and fixed at regular intervals to analyze the localization of TDP-43 by immunofluorescence. Nuclear envelope and NPC integrity was monitored by verifying that nuclei restrict the entry of a 70 kD fluorescent dextran (**Suppl fig 3A**), since the permeability barrier of the NPC increasingly excludes cargoes above 30-60 kD (Mohr *et al*, 2009; Timney *et al*, 2016). Using a luciferase-based reporter assay, we analyzed ATP levels following permeabilization and found that the total ATP concentration drops by ~97% immediately after digitonin treatment, compared to intact HeLa cells (**Suppl fig 3B**), confirming low ATP levels in the permeabilized cell system.

**Figure 3.**
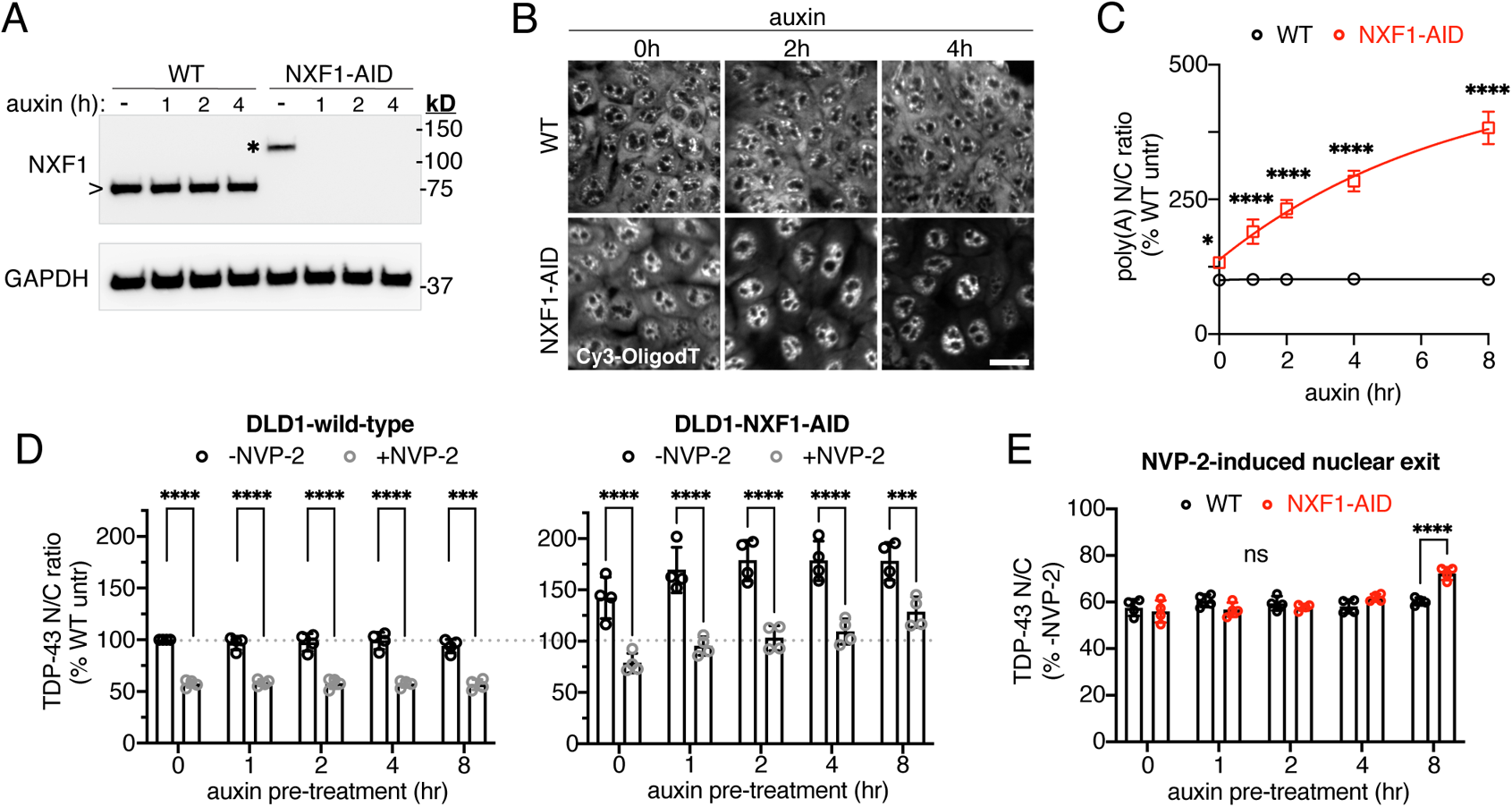
Acute NXF1 ablation does not alter NVP-2-induced TDP-43 nuclear export. A. NXF1 immunoblot in DLD1-wild-type vs. DLD1-NXF1-AID cells treated with 0.5 mM auxin for 0-4 h. Note the increased molecular weight for AID-tagged NXF1 (*) vs endogenous NXF1 (>). B. Poly(A)-FISH (Cy3-OligodT(45)) in DLD1-wild-type vs. DLD1-NXF1-AID cells treated with 0.5 mM auxin for designated time. Scale bar = 25 µm. C. Poly(A)-RNA N/C ratio in DLD1-wild-type vs. DLD1-NXF1-AID cells treated with 0.5 mM auxin for 0-8 h. Data are expressed as percent wild-type untreated cells. N = mean of 3476 cells/well in each of 4 independent biological replicates. Mean ± SD is shown. *p<0.05, ****p<0.0001 versus wild-type, by 2-way ANOVA with Sidak’s multiple comparisons test. D. TDP-43 N/C ratio in DLD1-wild-type cells (left) and DLD1-NXF1-AID cells (right) pretreated with 0.5 mM auxin for 0-8 h, followed by 2 h auxin only (-NVP-2) or auxin + 250 nM NVP-2 (+NVP-2). Data in both panels are normalized to untreated DLD1-wild-type cells (dotted line). E. TDP-43 N/C ratio in NVP-2 treated vs. untreated cells (+NVP-2 / −NVP-2) in DLD1-wild-type (black) vs. DLD1-NXF1-AID (red) cells. These are the same data as in (D), adjusted for differences in the steady-state TDP-43 N/C ratio, to permit comparison of NVP-2-induced TDP-43 nuclear exit. In D-E, N = mean of 4610 cells/well in each of 4 independent biological replicates. Mean ± SD is shown. ****P<0.0001 for indicated comparisons by 2-way ANOVA with Sidak’s multiple comparisons test.

Next, we identified conditions permissive of TDP-43 nuclear export. Upon plasma membrane permeabilization and maintenance in buffer, with no added transport receptors or ATP, we saw a progressive decrease in nuclear TDP-43 immunofluorescence, to about 50% after 1 h, in cells heated to a physiological temperature at 37°C (**Fig. 2B-C**). However, there was only minimal change in nuclear TDP-43 over an identical time period when cells were permeabilized and kept at 4°C. These results indicate that, while TDP-43 can passively leave the nucleus in low-ATP conditions at 37°C, temperature-sensitive mechanism(s) hinder TDP-43 nuclear exit at 4°C beyond the expected ~linear reduction of free diffusion with reduced temperature (Soh *et al*, 2010). To further test the passive nature of the observed export, we added 1,6-hexanediol, an aliphatic alcohol that reversibly disrupts the phenylalanine-glycine (FG)-permeability barrier lining the central channel of the NPC (Ribbeck & Görlich, 2002). 1,6-hexanediol caused a marked, dose-dependent acceleration of TDP-43 nuclear efflux, suggesting that nuclear pore permeability is a rate-limiting factor in TDP-43 nuclear export (**Fig. 2D–E**). Interestingly, 1,6-hexanediol accelerated TDP-43 nuclear exit not only at 37°C but also at 4°C, suggesting that the FG-repeat permeability barrier is temperature-sensitive and contributes, at least in part, to the hindrance of TDP-43 passive export at 4°C. Of note, 1,6-hexanediol also reversibly inhibits TDP-43 liquid-liquid phase separation (LLPS) over the concentration range used (Gopal *et al*, 2017; Mann *et al*, 2019). Dissolution of nuclear TDP-43 phase condensates could also contribute to the acceleration of TDP-43 nuclear export if LLPS affects the availability of TDP-43 monomers for diffusion across the NPC, which remains to be investigated. Consistent with lack of involvement of XPO1 in TDP-43 export, addition of the XPO1 inhibitor leptomycin B (LMB, 100 nM) during the permeabilization, washing, and export phases did not alter TDP-43 nuclear efflux (**Suppl fig 3C**). Next, we tested the effect of adding ATP and saw no change in the rate of nuclear export at concentrations far exceeding those that support active nuclear transport in permeabilized cell systems (Cassany & Gerace, 2009; Hayes *et al*, 2020, 2021) (**Suppl fig 3D**). To confirm that the data are not confounded by TDP-43 import back into the nucleus, we tested the ability of our permeabilized cell system to support active nuclear import. Although no import receptors or Ran cycle proteins were added to the assay, residual importins do remain at NPCs upon cell permeabilization (Kapinos *et al*, 2017). Therefore, we analyzed the capacity for nuclear import of Rango, a FRET sensor and direct importin β cargo (Kalab *et al*, 2006; Hayes *et al*, 2020), and saw no import in our lysate-free conditions, with or without added ATP (**Suppl fig 3E**). Our findings are therefore unlikely to be confounded by the nuclear re-entry of TDP-43.

Taken together, these data demonstrate that TDP-43 nuclear efflux in permeabilized cells is accelerated by heat and 1,6-hexanediol, but not ATP, consistent with nuclear export by passive diffusion through NPC channels.

### TDP-43 export is independent of NXF1-mediated mRNA export

The observation that TDP-43 exits the nucleus in permeabilized cells under passive conditions, and its exit is not accelerated by ATP, suggests that canonical, energy-requiring RNA export pathways are unlikely to be required for TDP-43 nuclear export. This includes the exportin-family of receptors which rely on the Ran-GTPase gradient (e.g. XPO1, XPOT, and XPO5), and the TREX bulk mRNA export pathway, which is independent of Ran but requires the activity of ATP-dependent RNA helicases, including UAP56 and DDX19b, for mRNP export complex assembly and disassembly (Okamura *et al*, 2015). In support of this, individual siRNA knockdown of XPO1, XPO5, XPOT, the NXF1 mRNA export receptor, and AlyREF, a TREX complex protein that participates in NXF1-recruitment, did not alter TDP-43 steady-state localization (Archbold *et al*, 2018; Ederle *et al*, 2018). Similarly, TDP-43 RRM mutations did not prevent nuclear export in the heterokaryon assay (Ederle *et al*, 2018). However, XPO1, XPO7, and NXF1 overexpression promoted TDP-43 cytoplasmic redistribution (Archbold *et al*, 2018). Of note, NXF1 has also been identified as a potential TDP-43 interacting protein (Freibaum *et al*, 2010).

To further exclude a role for the NXF1/TREX pathway in TDP-43 nuclear export, we utilized a DLD1 cell line in which an auxin-inducible degron (AID) tag was introduced into the endogenous NXF1 locus via CRISPR (Aksenova *et al*, 2020). TIR1 ligase, which drives ubiquitination of AID-tagged proteins upon auxin-mediated recruitment, was separately integrated at the C-terminus of the nuclear protein RCC1 via a self-cleavable P2A sequence. This enables rapid and complete degradation of the NXF1 protein within 1 h of exposure to 0.5 mM auxin (**Fig 3A**). Due to TIR1 leakage, DLD1-NXF1-AID cells showed mildly reduced NXF1 expression even prior to auxin administration (**Fig 3A** asterisk). To examine the effect of NXF1 ablation on mRNA export, DLD1-wild-type and DLD1-NXF1-AID cells were treated with auxin for 0-8 h, fixed, and poly(A)-RNA N/C localization was analyzed by fluorescence in situ hybridization (FISH) (**Fig 3B-C**). DLD1-NXF1-AID cells showed an ~30% increase in the poly(A)-RNA N/C ratio even prior to auxin administration (p<0.05 vs. DLD1-wild-type), consistent with the reduced NXF1 expression. Auxin induced further, progressive accumulation of nuclear poly(A)-RNA to >300% of baseline by 8 h, consistent with the inhibition of mRNA export (**Fig 3B-C**). To determine the effect of NXF1 ablation on transcriptional blockade-induced TDP-43 nuclear export, cells were treated with auxin for 0-8 h, followed by a 2 h incubation ± NVP-2 prior to fixation and TDP-43 immunostaining (**Fig 3D-E**). Interestingly, DLD1-NXF1-AID cells showed an increased steady-state TDP-43 N/C ratio compared to DLD1-wild-type cells that increased with auxin treatment, presumably due to TDP-43 binding to increased nuclear poly(A)-RNA (**Fig 3D**).

However, after adjusting for this increased steady-state N/C ratio, there was no delay in NVP-2-induced TDP-43 nuclear export in DLD1-NXF1-AID cells until 8h, when subtle slowing was observed (**Fig 3E**). These results suggest that inhibiting mRNA export promotes TDP-43 nuclear accumulation, but the NXF1 receptor itself does not mediate TDP-43 export across the NPC.

### TDP-43 nuclear localization depends on binding to GU-rich RNAs

Since TDP-43 is primarily nuclear, despite its ability to diffuse through the NPC, we reasoned that TDP-43 localization must depend on intranuclear sequestration, perhaps binding to RNA given the rapid nuclear efflux induced by transcriptional inhibition and nuclear accumulation conferred by mRNA export blockade. To test this hypothesis, we added increasing concentrations of RNase A to permeabilized cells and monitored the effect on TDP-43 nuclear localization (**Fig 4A-C**). In contrast to untreated controls, RNase treatment caused rapid TDP-43 nuclear efflux over 30 minutes even at 4°C, suggesting that RNA indeed tethers TDP-43 within the nucleus. Interestingly, following RNase treatment, residual endogenous nuclear TDP-43 formed puncta (**Fig 4B arrows**), as reported following nuclear RNase microinjection in living cells (Maharana *et al*, 2018), and similar to the expression pattern of tagged, overexpression constructs of TDP-43 RNA binding mutants (Ayala *et al*, 2008; Elden *et al*, 2010; Wang *et al*, 2020; Yu *et al*, 2020).

**Figure 4.**
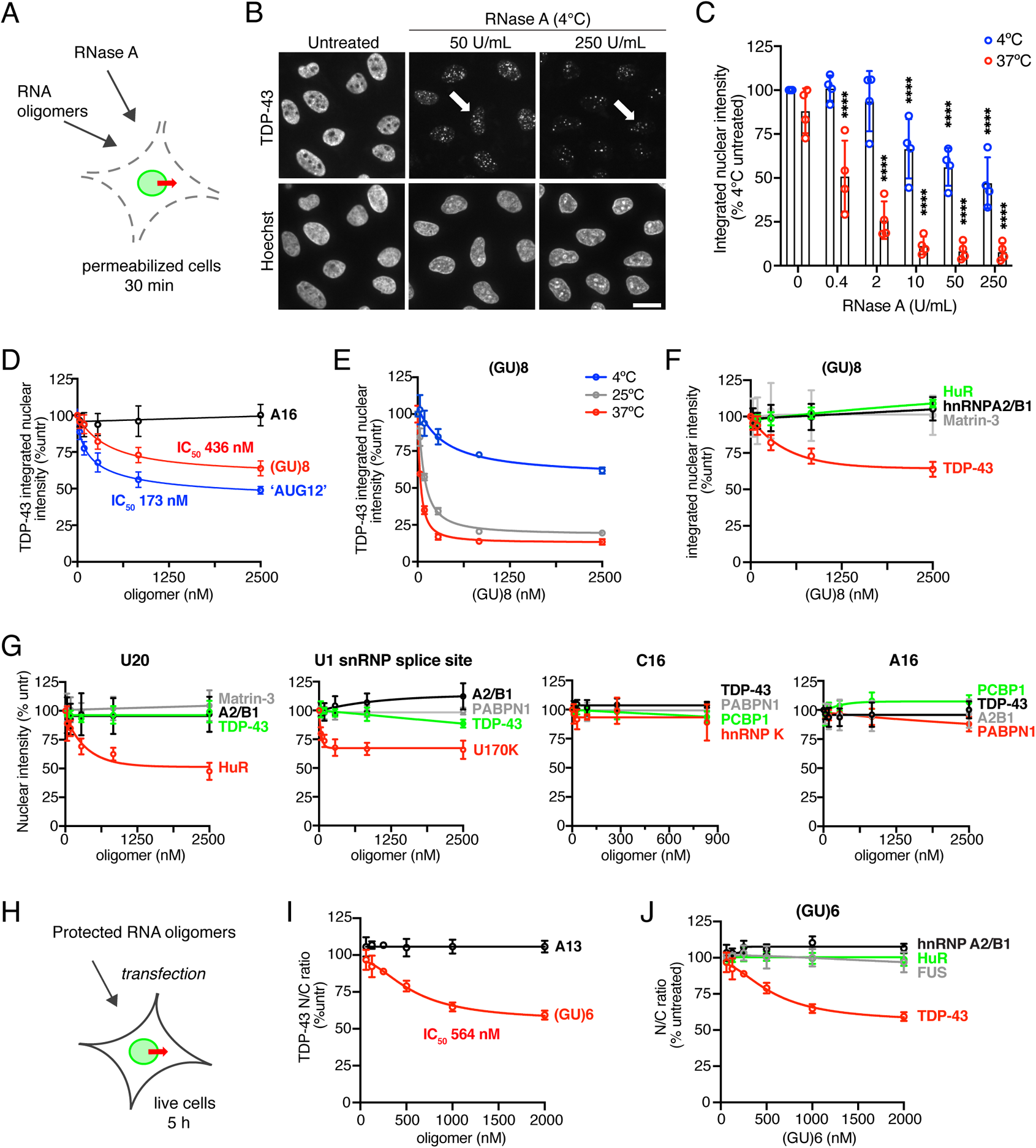
RNase and GU-rich oligomers induce TDP-43 nuclear efflux. A. Schematic depicting addition of RNase (A-C) or RNA oligomers (D-G) to permeabilized cell TDP-43 passive export assay. B. Endogenous TDP-43 immunofluorescence (top) and Hoechst (bottom) in permeabilized HeLa cells incubated for 30 min at 4°C with or without RNase A. Arrows highlight formation of nuclear puncta by residual nuclear TDP-43. Scale bar = 25 μm. C. TDP-43 integrated nuclear intensity in permeabilized HeLa cells incubated for 30 min at 4°C vs. 37°C with increasing concentrations of RNase A. Data are expressed as percent untreated cells at 4°C. N = mean of 2552 cells/well/condition in each of 4 independent biological replicates. ****p<0.0001 versus time 0 by 2-way ANOVA with Tukey’s multiple comparisons test. D. TDP-43 integrated nuclear intensity in permeabilized HeLa cells incubated for 30 min at 4°C with increasing concentrations of A16, (GU)8, or ‘AUG12’ oligomers. Data are expressed as percent untreated cells. N = mean of 2968 cells/well/condition in each of 4 independent biological replicates. E. TDP-43 integrated nuclear intensity in permeabilized HeLa cells incubated at 4°C, 25°C or 37°C for 30 min with increasing concentrations of (GU)8. Data are expressed as percent untreated cells. N = mean of 1039 cells/well/condition in each of 2 technical replicates. F. Integrated nuclear intensity of designated nuclear RBPs in permeabilized HeLa cells incubated for 30 minutes at 4°C with increasing concentrations (GU)8. Data are expressed as percent untreated cells. Note: TDP-43 data are the same as in (D). N = mean of 3421 cells/well in each of 4 independent biological replicates. G. Integrated nuclear intensity of designated nuclear RBPs in permeabilized HeLa cells incubated for 30 minutes at 4°C with increasing concentrations of the indicated RNA oligomers. Data are expressed as percent untreated cells. N = mean of 3293 cells/well/condition in each of 3-4 independent biological replicates. H. Schematic of live HeLa cell transfection with protected RNA oligomers. I. TDP-43 N/C ratio 5 h after transfection with protected A13 or (GU)6 RNA oligomers. Data are expressed as percent untreated cells. N = mean of 2887 cells/well/condition in each of 3 independent biological replicates. J. RBP N/C ratios 5 h after transfection with protected (GU)6 RNA oligomers. Data are expressed as percent untreated cells. Note: TDP-43 data are the same as those shown for (GU)6 in (I). N = mean of 2555 cells/well/condition in each of 3 independent biological replicates. In F,G,J: The RBP labeled in red corresponds to the most closely-predicted binding partner for that motif, and green indicates an RBP with a moderately-similar motif. Gray and black labels correspond to RBPs with no or low predicted binding activity to the given sequence. In D-J: RNA oligomer sequences are detailed in **Table 2**. Mean ± SD is shown. IC50 values were calculated by non-linear regression.

**Table 2.**
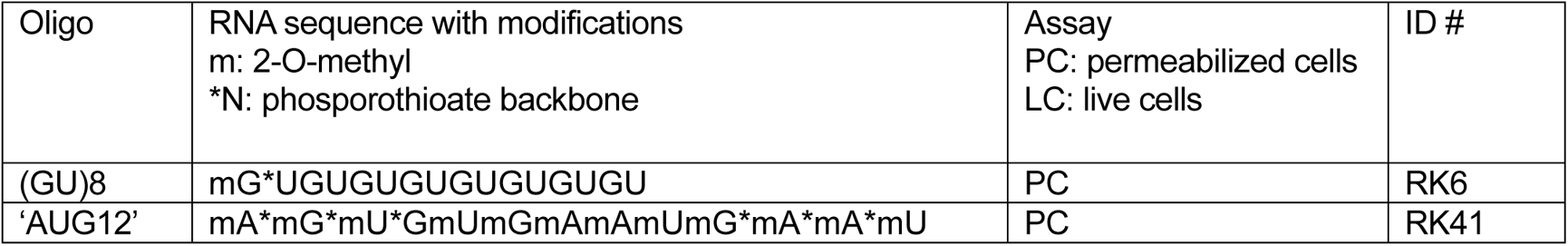
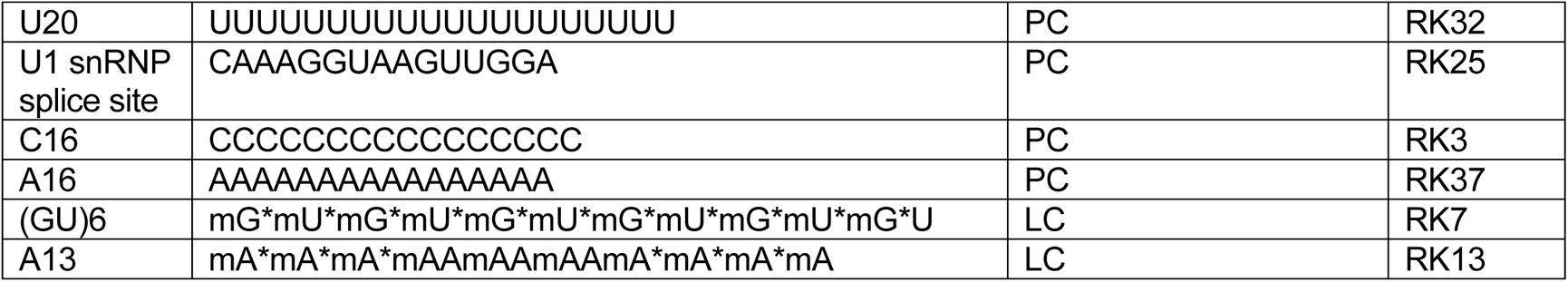
RNA oligonucleotides.

Furthermore, in addition to the temperature-induced changes in the NPC permeability barrier discussed above (**Fig. 2D-E**), these results suggest that temperature-sensitive dissociation from RNAs contributes to the drastically reduced passive TDP-43 nuclear exit at 4°C versus 37°C. To assess nuclear integrity, we also analyzed the effect of RNase on the nuclear localization of DDX19b (54 kD) and Nup50 (50 kD), two proteins of similar size to TDP-43 that are localized to the NPC and nucleoplasm (**Suppl fig 4A**). The nuclear intensity of both proteins was unaffected at 4°C, and modestly diminished at 37°C, but overall remarkably stable compared to the rapid nuclear efflux observed for TDP-43. The observation of the critical role of RNA in TDP-43 nuclear localization prompted us to test whether RNA degradation post-permeailization contributes to the TDP-43 nuclear efflux we observed in our passive export assay (**Suppl fig 4B**). We saw no difference in the rate of TDP-43 nuclear efflux in permeabilized cells kept at 37°C for 30 or 60 min, with or without RNasin, suggesting that RNA degradation does not contribute to the passive export we observe. Nevertheless, RNasin was added to the transport buffer for all permeabilized cell assays as a precautionary measure.

CLIP-seq studies demonstrate that TDP-43 preferentially binds repetitive GU-rich sequences, particularly within introns (Polymenidou *et al*, 2011; Tollervey *et al*, 2011), and the RRM domains optimally recognize a synthetic ‘AUG12’ (GUGUGAAUGAAU) motif (Lukavsky *et al*, 2013). To investigate the specificity of the RNA-mediated nuclear tethering of TDP-43, we added synthetic RNA oligomers designed as ‘decoys’ mimicking the preferred GU-rich binding sites to the permeabilized cell passive nuclear export system (**Fig 4D-G**). Following permeabilization, increasing concentrations of (GU)8, ‘AUG12’, or A16 oligomers were introduced, and cells incubated at 4°C for 30 min (**Fig 4D**). Both ‘AUG12’ and (GU)8 induced dose-dependent nuclear efflux of TDP-43, presumably by competitive dissociation of TDP-43 from endogenous nuclear RNAs, freeing TDP-43 to passively exit the nucleus. The A16 control oligomer had no effect.

Incubating permeabilized cells with (GU)8 at increasing temperatures (25°C and 37°C) markedly accelerated (GU)8-induced TDP-43 export (**Fig 4E**), consistent with the expected temperature-dependency of TDP-43 free diffusion, NPC permeability, and temperature-sensitive dissociation of TDP-43 from nuclear RNAs.

Next, we evaluated the selectivity of the oligomer-induced nuclear efflux comparing different RNA motifs and RBPs in the permeabilized cell system. (GU)8 induced nuclear efflux of TDP-43 but not three other RBPs, including HuR which preferentially recognizes related U-rich motifs (**Fig 4F**) (e.g. UUGGUUU, http://rbpmap.technion.ac.il/ (Paz *et al*, 2014; Silanes *et al*, 2004; Lebedeva *et al*, 2011)). Conversely, polyU (U20) induced the nuclear efflux of HuR but not TDP-43 or other RBPs (**Fig 4G**). An oligomer based on the U1 snRNP splice site sequence (CAAAGGUAAGUUGGA (Kondo *et al*, 2015)) selectively induced the nuclear export of the U1 small nuclear ribonucleoprotein 70 kDa (U1-70K or snRNP70), with no significant effect on the localization of TDP-43, hnRNPA2/B1, or PABPN1. C16 and A16 oligomers failed to induce nuclear export of any RBPs tested, including those predicted to bind (i.e. hnRNPK and PABPN1, respectively (Choi *et al*, 2009; Goss & Kleiman, 2013)). Thus, at least in the permeabilized cell system, these results show that nuclear localization of a subset of RBPs, including TDP-43, depends on remarkably specific and selective binding to their preferred binding motifs within nuclear RNAs.

Finally, we tested the ability of RNA oligomers to alter TDP-43 nuclear localization in live cells (**Fig 4H**). HeLa cells were transfected with increasing concentrations of (GU)6 or A13 oligomers that were modified by the addition of 2’-O-methyl groups and phosphorothioate bonds to prevent degradation by cellular RNases. Cells were fixed 5 h after transfection for immunostaining and analysis. As in permeabilized cells, (GU)6 but not A13 induced dose-dependent TDP-43 nuclear efflux (**Fig 4I**). Again, this was specific for TDP-43 as there was no change in the N/C ratio of HuR, FUS, or hnRNPA2/B1 (**Fig 4J**). Together, these findings confirm that in live cells, specific binding to endogenous GU-rich nuclear RNAs opposes the tendency of TDP-43 to passively leave the nucleus.

### Inhibition of pre-mRNA splicing promotes TDP-43 nuclear accumulation

Since GU-rich nuclear RNA binding appears critical for TDP-43 nuclear retention, we hypothesized that TDP-43 nuclear accumulation relies on the abundance of GU-rich intronic sequences in newly-transcribed pre-mRNAs prior to splicing and degradation of released introns. Since inhibition of pre-mRNA splicing stalls intron excision and leads to nuclear accumulation of unspliced pre-mRNAs (Carvalho *et al*, 2017), we used two splicing inhibitors, the Ginkgo biloba tree-derived bioflavinoid isoginkgetin (IGK) (O’Brien *et al*, 2008) and the bacterial macrolide pladienolide B (PLB) (Sato *et al*, 2014), to test whether nuclear intron accumulation could reduce TDP-43 nuclear efflux. PLB binds and inhibits SF3B1 in the U2 snRNP in the first step of spliceosome assembly (Kotake *et al*, 2007). The precise target of IGK has not yet been identified, but it blocks the complex A to complex B transition during spliceosome assembly, downstream of PLB (O’Brien *et al*, 2008). After a 4 h exposure, both IGK and PLB induced accumulation of introns as determined by qRT-PCR analysis with primers targeting the exon/intron junction of selected housekeeping genes, normalized to the expression of U6 snRNA, an unspliced transcript of RNA polymerase III (O’Brien *et al*, 2008) (**Suppl fig 5**). Over the same time course, there was a dose-dependent increase in the steady-state N/C ratio of TDP-43 (**Fig 5A-B, D-E**). Remarkably, when cells were incubated for 4 h with IGK or PLB, and NVP-2 was added to induce transcriptional blockade and TDP-43 nuclear efflux, both IGK and PLB promoted TDP-43 nuclear retention, in a dose-dependent manner (**Fig 5C,F**). This effect was more potent for IGK than PLB, albeit at higher doses (IGK IC_50_ = 62 µM, PLB IC_50_ = 6 nM). Together with the observed effects of GU-rich oligomers in permeabilized and live cells, these data strongly suggest that TDP-43 nuclear localization depends on the abundance of its nuclear intronic pre-mRNA binding sites.

**Figure 5.**
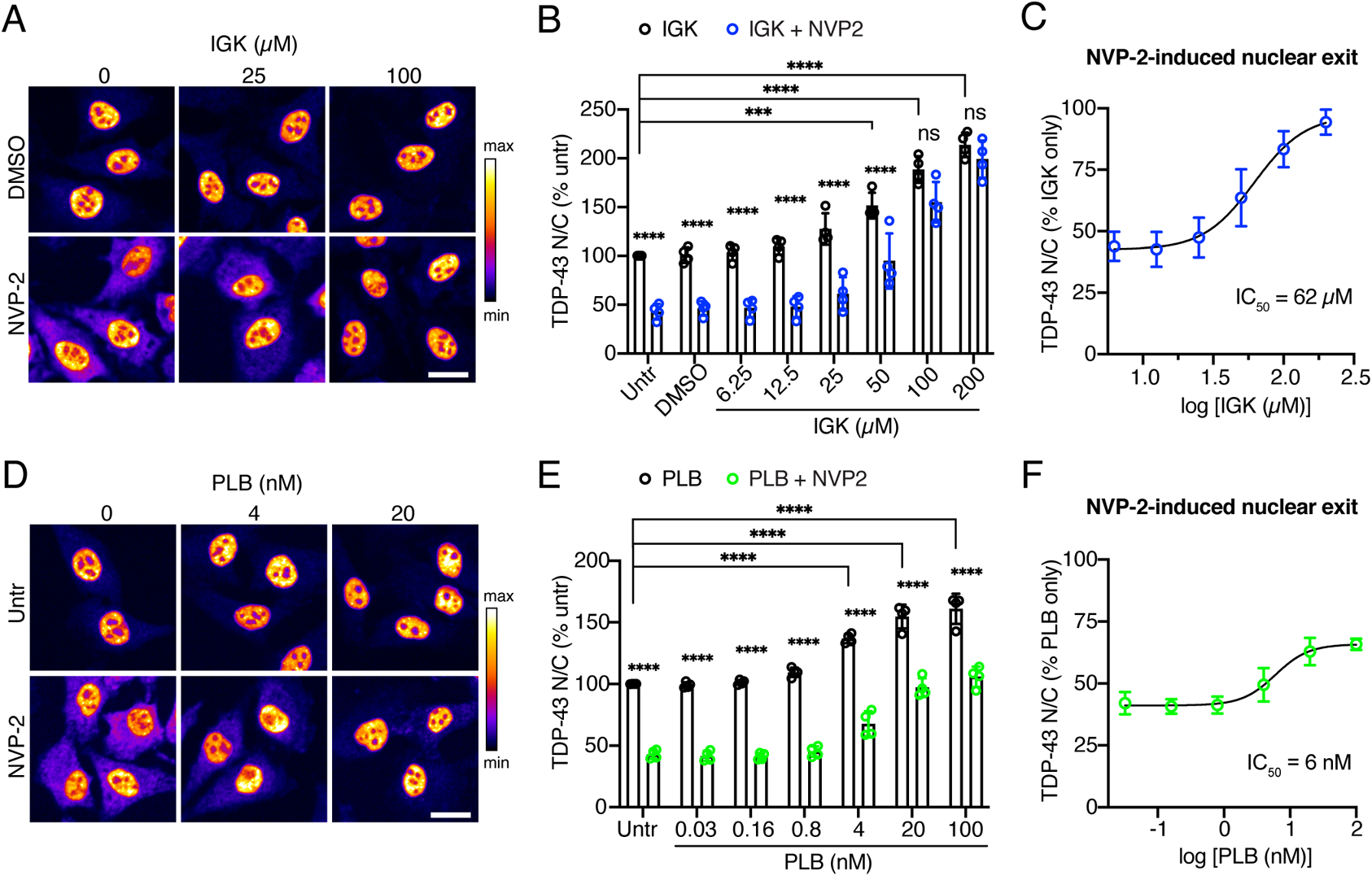
Inhibition of pre-mRNA splicing promotes TDP-43 nuclear accumulation. A. TDP-43 immunofluorescence in HeLa cells treated with IGK for 4 h, followed by 2 h of IGK only or IGK + 250 mM NVP-2. Scale bar = 25 µm. B. TDP-43 N/C in IGK or IGK + NVP-2-treated cells, expressed as % untreated. C. TDP-43 N/C ratio (same data as part B) expressed as % IGK only, to permit comparison of NVP-2-induced nuclear exit. IC_50_ calculated by non-linear regression. D. TDP-43 immunofluorescence in HeLa cells treated with PLB for 4 h, followed by 2 h of PLB only or PLB + 250 mM NVP-2. Scale bar = 25 µm. E. TDP-43 N/C in PLB or PLB + NVP-2-treated cells, expressed as % untreated. F. TDP-43 N/C ratio (same data as part E) expressed as % PLB only, to permit comparison of NVP-2-induced nuclear exit. IC_50_ calculated by non-linear regression. In A,C: The intensity histogram for each image was independently maximized across the full range to enable comparison of nuclear vs. cytoplasmic signal, and a pseudo-color LUT was then applied (see linear scale at right). In B,C: N= mean of 2357 cells/well in each of 4 independent biological replicates. Mean ± SD is shown. In E,F: N= mean of 2720 cells/well in each of 4 independent biological replicates. Mean ± SD is shown. In B,C,E,F: ***p<0.001, ****p<0.0001 by 2-way ANOVA with Tukey’s multiple comparisons test. Comparisons as indicated with untreated cells (brackets) or between NVP-2-treated and untreated cells at each dose of splicing inhibitor (no brackets). See **Suppl fig 5** for validation of IGK and PLB-induced intron accumulation by qRT-PCR.

### TDP-43 RRM domains confer RNA-dependent TDP-43 nuclear localization

The RNA-binding properties of the TDP-43 RRM domains have been extensively characterized (Buratti & Baralle, 2001; Lukavsky *et al*, 2013; Cohen *et al*, 2015; Flores *et al*, 2019). Although mutation or deletion of the RRM domains does not abolish TDP-43 nuclear localization (Elden *et al*, 2010; Yu *et al*, 2020), these observations arise from fluorescently tagged, high molecular weight TDP-43 constructs, which markedly inhibits TDP-43 nuclear export (**Suppl fig 1K**; Ederle *et al*, 2018; Pinarbasi *et al*, 2018). However, Ayala and colleagues previously generated TDP-43 RRM mutants with a small (~1 kD) FLAG tag and demonstrated increased cytoplasmic localization by N/C fractionation and immunoblotting (Ayala *et al*, 2008). To further assess the role of the RRM domains for TDP-43 nuclear localization and export without markedly increasing its molecular weight, we generated a series of constructs expressing V5-tagged wild-type and RRM mutants of TDP-43 (**Fig 6A**), including mutations of phenylalanine residues (5F◊L) and acetylation sites (2K→Q) that are critical for RNA binding (Buratti & Baralle, 2001; Cohen *et al*, 2015), and a complete RRM deletion (ΔRRM1-2). Constructs were transiently transfected into a stable HeLa cell line largely depleted of TDP-43 by CRISPR (**Suppl fig 6A-B**) (generously provided by S. Ferguson (Roczniak-Ferguson & Ferguson, 2019)) to avoid potential confounding effects of hetero-oligomerization with endogenous TDP-43. High-content image analysis showed a significant drop in the steady-state N/C ratio of all three RRM mutants, presumably exaggerated for ΔRRM1-2 due to its small size (29 kD) in comparison to the point mutants (45 kD) (**Fig 6A-B**). In addition to reduced nuclear accumulation of the RRM mutants, shuttling in response to NVP-2 was abolished. Next, we confirmed these findings with N/C fractionation and immunoblotting (**Fig 6C-D**). The immunoblotting results corroborated our image-based analyses, showing a marked reduction in the N/C ratio of TDP-43 in all three RRM mutants and no response to NVP-2. Thus, TDP-43 RRM domains are required for nuclear TDP-43 localization and RNA-regulated shuttling behavior.

**Figure 6.**
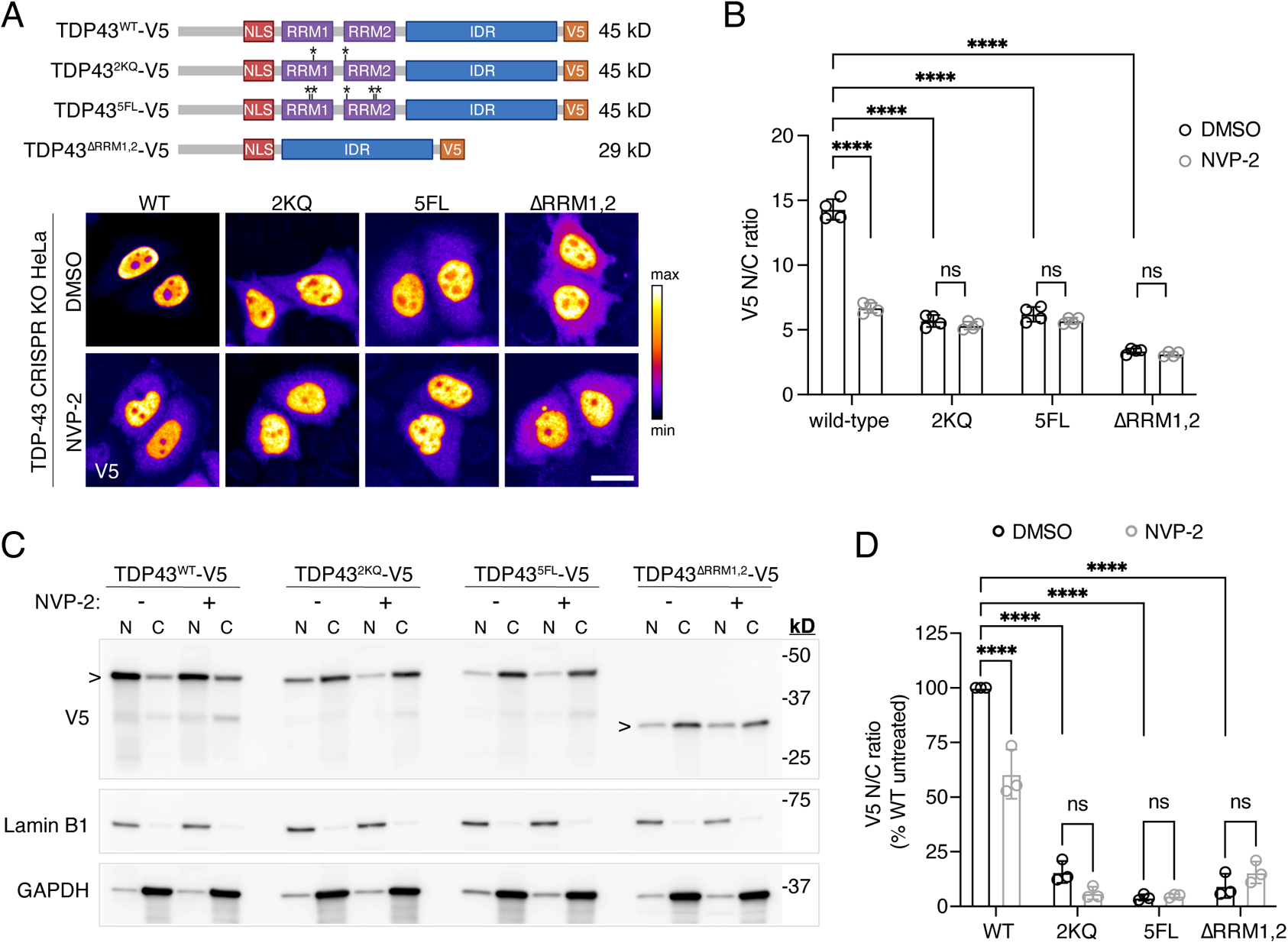
Mutation or deletion of TDP-43 RRM domains disrupts TDP-43 nuclear localization. A. Schematic of V5-tagged TDP-43 RRM mutant constructs (top) and V5 immunofluorescence (bottom) in transiently transfected TDP-43 CRISPR KO HeLa cells, after 1 h treatment with DMSO vs. 250 nM NVP-2. The intensity histogram for each image was independently maximized across the full range to enable comparison of nuclear vs. cytoplasmic signal, and a pseudo-color LUT was then applied (see linear scale at right). See **Suppl fig 6** for validation of CRISPR knockdown of endogenous TDP-43. Scale bar = 25 µm. B. Raw N/C ratio of TDP-43 wild-type vs. RRM mutant constructs in transiently transfected TDP-43 CRISPR KO HeLa cells, following 1 h treatment with DMSO or 250 nM NVP-2. N= mean of 1015 cells/well in each of 4 replicates (2 technical and 2 biological). Mean ± SD is shown. C. N/C fractionation and immunoblotting of TDP-43 CRISPR KO HeLa cells, transiently transfected with V5-tagged TDP-43 constructs and treated for 3 h with DMSO vs. 250 nM NVP-2. Lamin B1 = nuclear marker. GAPDH = cytoplasmic marker. D. N/C ratio of V5 nuclear vs. cytoplasmic signal calculated from N=3 immunoblots (derived from 2 biological replicates). Ratios are expressed as % WT DMSO-treated cells. In B,D: ns = not significant, ****p<0.0001 by 2-way ANOVA with Tukey’s multiple comparisons test.

Of note, coalescence of the V5-tagged RRM mutants into nuclear puncta was only rarely observed, compared to widespread puncta observed in cells transfected with YFP-tagged TDP-43-5F→L (**Suppl fig 6C,F**) (Elden *et al*, 2010). Since 27kD-YFP contains the same three residues (F223, L221, A206) known to cause dimerization of GFP (Day & Davidson, 2009), the presumed nuclear phase separation behavior of TDP-43 RRM mutants could be exacerbated by the size or dimerization activity of GFP-derived fluorophores that can drive aberrant localization of tagged proteins in live cells (Snapp *et al*, 2003; Falcón-Pérez *et al*, 2005). Indeed, we also saw a trend toward increased nuclear puncta formation by wild type TDP-43-YFP compared to TDP-43-V5, though never to the extent of TDP-43-5FL-YFP (**Suppl fig 6F**).

### Role of HSP70 chaperone activity in nuclear/cytoplasmic localization of RNA-free TDP-43

The HSP70 family of chaperones was recently shown to regulate the nuclear phase separation of RNA-free TDP-43 into droplets or “anisosomes” (Yu *et al*, 2020). We wondered whether, in addition to NLS-mediated nuclear import, HSP70 activity might therefore contribute to the nuclear localization of TDP-43 RRM mutants. First, we confirmed that HSP70 inhibition strongly modifies fluorescent nuclear puncta in TDP-43 CRISPR KO cells transiently transfected with TDP-43-YFP or TDP-43-5FL-YFP (**Suppl fig 6C**). TDP-43-5FL-YFP-expressing cells exhibited striking and widespread nuclear puncta, observed in 99.8% of cells vs. 35% of wild-type TDP-43-YFP-expressing cells (**Suppl fig 6F**). Consistent with the report of Yu and colleagues, treatment with the HSP70 small molecule inhibitor VER 155008 (HSP70i, 50 µM) (Massey *et al*, 2010; Yu *et al*, 2020) induced a steady decline in the number of TDP-43-5FL-YFP puncta per nucleus (**Suppl fig 6D**), whereas no change was seen in TDP-43-YFP-expressing cells. Since the YFP tag alters nuclear localization and export (**Suppl fig 1K**), next we verified that HSP70 also regulates nuclear puncta formation using our V5-tagged RRM mutants (**Suppl fig 6E**). As shown in **Fig 6**, the V5-tagged proteins formed far fewer nuclear puncta, observed in 9% of WT-V5, 17% of 2KQ-V5, 14% of 5FL-V5, and 13% of ΔRRM1,2-V5-transfected cells (**Suppl fig 6F**). Puncta consisted of fine granules in 2KQ-V5 and 5FL-V5-expressing cells and larger spherical/shelled structures in the ΔRRM1,2-V5-transfected cells (arrows in **Suppl fig 6E**), and were more likely to be observed in higher-expressing cells across all constructs.

HSP70i induced a significant increase in the number of cells with nuclear puncta in both 2KQ-V5 and ΔRRM1,2-V5-transfected cells, and a similar trend in 5FL-V5-transfected cells (**Suppl fig 6E-F**), accompanied by increased mean puncta/nucleus and mean puncta area that was never observed in WT-V5-expressing cells (**Suppl fig 6H**). Thus, while HSP70i induced the simplification of pre-existing nuclear puncta in TDP43-5FL-YFP-expressing cells, it induced the formation of new and more numerous nuclear puncta in cells expressing V5-RRM mutants with a smaller, dimerization-free tag. Given reduced antibody penetration into densely-packed liquid spherical shells of larger anisosomes (Yu *et al*, 2020), it is possible that the V5 immunofluorescence (IF) under-represents the amount of V5-TDP-43 residing in nuclear puncta. However, N/C fractionation and immunoblotting (using denaturing conditions) showed a remarkably similar distribution of V5-tagged TDP-43 to the results obtained with IF (**Fig 6**), indicating that only a small fraction of the total V5-TDP-43 is enclosed in structures invisible by IF. Overall, HSP70i treatment is clearly altering the nuclear distribution of V5-tagged RRM mutants (**Suppl fig 6**), confirming potent chaperone activity in regulating the sub-nuclear localization of RNA-free TDP-43 in our model system.

Next, we analyzed the effect of HSP70i on the nuclear/cytoplasmic partitioning of TDP-43-V5 versus V5-tagged RRM mutants at steady state and following NVP-2-induced transcriptional blockade (**Suppl fig 6I**). Again, NVP-2 induced cytoplasmic shuttling of wild-type TDP-43-V5 and had no effect on the RRM mutants. Interestingly, HSP70i did not further compromise the ability of RRM-mutant TDP-43 to remain in the nucleus. Rather, HSP70i induced a modest (10-15%) increase in the TDP-43 N/C ratio of all three RRM mutants that was not observed for wild-type TDP-43-V5. Thus, HSP70-dependent nuclear chaperone activity appears to modestly favor the nuclear export of RNA-free TDP-43, by an unknown mechanism.

These data suggest that HSP70 chaperones do not contribute to the nuclear accumulation of RRM-mutant TDP-43, which is most likely attributable to ongoing active and passive nuclear import.

## Discussion

In this study, we investigated the mechanism of TDP-43 nuclear export and the regulatory role of RNA in maintaining TDP-43 nuclear localization. In permeabilized cells, TDP-43 readily exited the nucleus in low-ATP conditions, consistent with passive diffusion through NPCs. Acute depletion of NXF1 did not alter TDP-43 nuclear export in live cells, further excluding active TDP-43 co-export with mRNA. Three lines of evidence indicate that binding to GU-rich nuclear RNAs sequesters TDP-43 in nuclei and controls its availability for passive nuclear exit. (1) Degradation of nuclear RNAs by RNase treatment in permeabilized cells induced rapid TDP-43 nuclear efflux, demonstrating that RNA is required for TDP-43 nuclear sequestration. (2) In permeabilized and live cells, synthetic GU-rich oligomers induced nuclear TDP-43 exit, likely by competitive displacement of TDP-43 from endogenous GU-rich nuclear RNAs. Moreover, splicing inhibitors promoted TDP-43 nuclear accumulation and resistance to nuclear efflux upon transcriptional blockade, further supporting the critical role of TDP-43 binding to intronic pre-mRNAs for maintenance of nuclear localization. (3) Mutation or deletion of TDP-43 RRM domains strongly reduced TDP-43 nuclear enrichment and abolished its RNA-regulated shuttling. Taken together, these findings support a model (**Fig 7**) in which TDP-43 nuclear/cytoplasmic distribution results from a balance between active and passive TDP-43 nuclear import, nuclear sequestration by binding to GU-rich intronic pre-mRNAs, and passive nuclear export. In this model, TDP-43 moves in and out of the nucleus via a reaction-diffusion-controlled mechanism (Bastiaens *et al*, 2006; Soh *et al*, 2010) whereby transient formation of high molecular weight TDP-43 complexes with nuclear pre-mRNAs (‘reaction’) locally sequesters TDP-43 within nuclei and hinders availability of free TDP-43 for passive diffusion through NPC channels. Since inhibition of nuclear RNA synthesis and competitive displacement with GU-RNA ‘decoys’ both induced net TDP-43 nuclear exit in living cells, the active import rate of TDP-43 is clearly insufficient to balance passive export resulting from loss of nuclear pre-mRNA binding sites. Either a drop in nuclear pre-mRNA binding sites or mutation of the TDP-43 RRM domains steeply increases the nuclear abundance of TDP-43 that can passively diffuse through NPCs, leading to (near) equilibration of TDP-43 between the nucleus and cytoplasm. By maintaining the nuclear abundance of GU-rich intronic pre-mRNAs, the dynamic balance of transcription and pre-mRNA processing may ultimately be the primary upstream factor determining TDP-43 nuclear localization under physiological conditions.

**Figure 7.**
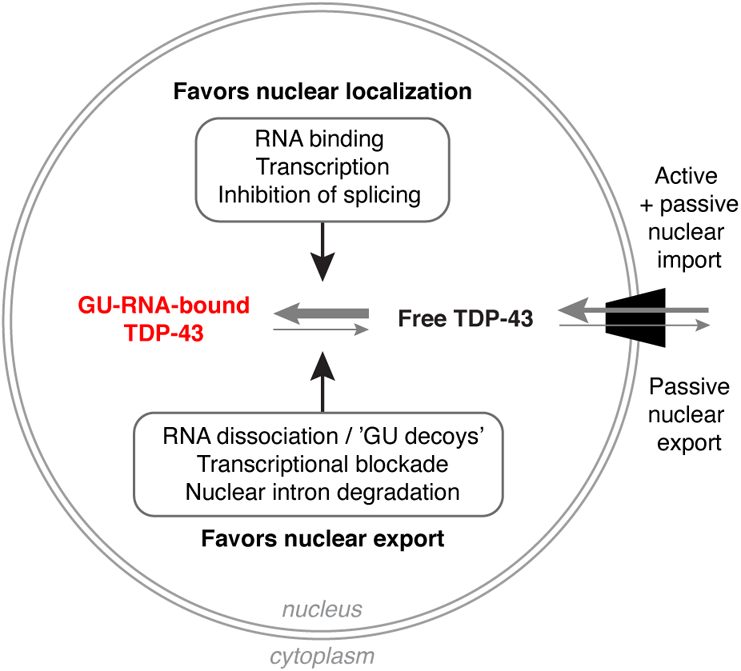
Mechanisms coupling TDP-43 nuclear localization with nuclear GU-rich pre-mRNA binding and abundance. Arrow size corresponds to predicted balance under normal physiologic conditions. See text for discussion.

### Transcriptional dependence of RBP nuclear enrichment

Consistent with previous studies (Piñol-Roma & Dreyfuss, 1992; Cáceres *et al*, 1998; Ayala *et al*, 2008; Ederle *et al*, 2018), we observed that transcriptional blockade altered the nuclear-cytoplasmic partitioning of numerous RBPs, ranging from prominent accumulation in the cytoplasm (TDP-43 and HuR) to a nearly two-fold nuclear increase (hnRNPC). Cytoplasmic-shifting RBPs included those with distinct RNA binding preferences, such as TDP-43 (Polymenidou *et al*, 2011; Tollervey *et al*, 2011; Lukavsky *et al*, 2013), HuR (Lebedeva *et al*, 2011), and hnRNPA1 (Bruun *et al*, 2016; Jain *et al*, 2017), as well as NXF1, which binds RNA in a sequence-independent manner (Tuck & Tollervey, 2013; Baejen *et al*, 2014). Similarly, RBPs retained or enriched in nuclei upon transcriptional inhibition included sequence-specific binders such as U2AF65 (Wu *et al*, 1999; Zorio & Blumenthal, 1999; Merendino *et al*, 1999) and the DEAD-box helicase DDX19b which has no sequence requirement (Lin *et al*, 2018). Thus, the diverse shuttling response of individual RBPs to perturbation of transcription appears to be defined by RBP-specific protein-RNA and protein-protein interactions rather than by a particular RNA motif.

Our mechanistic understanding of RNA synthesis-dependent N/C translocations of RBPs, including the factors mediating RBP nuclear export or retention, remains incomplete (Nakielny & Dreyfuss, 1999). Several RBP-specific functional domains have been identified to mediate nucleocytoplasmic shuttling, including the HuR nucleocytoplasmic shuttling (HNS) domain (Fan & Steitz, 1998), the K nuclear shuttling (KNS) motif of hnRNPK (Michael *et al*, 1997), the M9 signal required for transportin-mediated nuclear import and transcriptional blockade-induced export of hnRNPA1 (Michael *et al*, 1995), the nuclear retention signal of hnRNPC (Nakielny & Dreyfuss, 1996), and the SR domain of SF2 (SRSF1), which confers actinomycin D-dependent shuttling to other proteins (Cáceres *et al*, 1998). Here, we demonstrate that the TDP-43 RRM1-RRM2 domains mediate its transcriptional blockade-induced nuclear efflux and act as a nuclear retention domain via binding nuclear RNAs. Interestingly, a recent study showed that the depletion of cytoplasmic RNAs, via activation of the primarily cytoplasmic Xrn1 exonuclease-dependent mRNA decay, induced nuclear translocation of many RBPs, including the cytoplasmic poly(A) binding protein PABPC (Gilbertson *et al*, 2018). In a spatial inverse to our current model of TDP-43 shuttling, the findings of Gilbertson and colleagues suggest that the cytoplasmic abundance of poly(A)-RNAs may be responsible for maintaining cytoplasmic PABPC localization. Nucleocytoplasmic gradients of the preferred RNA binding motifs may therefore represent a broadly-applicable model for regulating RBP localization, which warrants further investigation.

### Passive diffusion of TDP-43 across the NPC

The FG permeability barrier of the NPC does not have a strict size cutoff; rather, diffusion of cargoes through the NPC is increasingly restricted over the 30-60 kD range (Mohr *et al*, 2009; Timney *et al*, 2016). Thus, 43 kD TDP-43 monomers are predicted to diffuse across the NPC with moderate efficiency. Indeed, we observed TDP-43 nuclear efflux in permeabilized cells in low-ATP conditions, that was accelerated by NPC permeabilization as expected for passive diffusion from the nucleus. These data corroborate predictions from live-cell assays showing marked size-restriction of TDP-43 nuclear export, including slowed TDP-43-tdTomato (+54 kD) export in the heterokaryon assay (Pinarbasi *et al*, 2018), lack of TDP-43-GCR_2_-EGFP_2_ (+119 kD) cytoplasmic recovery in a dexamethasone-induced shuttling assay (Ederle *et al*, 2018), and failure of TDP-43-YFP (+27 kD) nuclear efflux following transcriptional blockade (**Suppl fig 1K**). Delay in active nuclear import has been demonstrated for very large cargos (Paci *et al*, 2020), and the rate of active mRNP export is modestly size-dependent (Grünwald *et al*, 2011). However, the exquisite size-limitation of TDP-43 nuclear export in live-cell assays is most consistent with passive diffusion from the nucleus, as our permeabilized cell passive export assay confirms.

In addition to size, cargo surface properties also critically dictate NPC passage, including hydrophobic residues which augment NPC transport (Frey *et al*, 2018). Besides the glycine-rich C-terminal IDR, hydrophobic patches are present throughout the TDP-43 sequence (https://web.expasy.org/protscale/ (Gasteiger *et al*, 2005)), including within the N-terminal domain, RRM1, and RRM2, but their roles in NPC passage are not known. A recent study showed that intrinsically disordered proteins (up to ~63kDa) can passively transit through NPC channels (Junod *et al*, 2020), suggesting that the disordered state of the TDP-43 C-terminal IDR could potentially contribute to its passive nuclear exit. The potential role of TDP-43 hydrophobic and disordered regions for passive translocation through NPC channels remains to be examined.

### Nuclear–cytoplasmic GU-rich RNA gradients and TDP-43 shuttling

CLIP-seq studies have shown that TDP-43 preferably binds to long stretches of GU-rich intronic pre-mRNA sequences present in about 30% of genes (Polymenidou *et al*, 2011; Tollervey *et al*, 2011). Introns are in average much longer than exons in the predicted pre-mRNAs of human genes (7.5 kB vs 320 B (Lee & Rio, 2015)), and some are retained in exported mRNAs due to alternative splicing or incomplete debranching, and detectable in the cytoplasm as stable intron lariats (Talhouarne & Gall, 2018; Saini *et al*, 2019). However, a substantial proportion of nuclear intronic sequences, whether still contained in pre-mRNAs or excised, are likely short-lived, due to the tight coupling of transcription with pre-mRNA splicing, pre-mRNA quality surveillance, and rapid degradation via the nuclear exosome (Lee & Rio, 2015; Kilchert *et al*, 2016; Bresson & Tollervey, 2018). Active transcription could then potentially give rise to the N/C concentration gradient of the preferred TDP-43 binding sites in intronic GU-rich RNAs. The evident nuclear accumulation of newly-synthesized 5-EU-labeled RNAs in HeLa and neurons (**Fig 1A** and **Suppl fig 1A,F**) is indeed consistent with the existence of a steep N/C gradient of U-rich nascent RNAs. Recent rapid advances in CLIP-seq and related methods (Wheeler *et al*, 2018; Hafner *et al*, 2021) have yielded detailed insights into the RNA-RBP interactions and their dynamics (Nostrand *et al*, 2020). Despite this progress, currently available computational methods are not well suited for global quantification of particular sequence motifs (such as GU repeats) across all RNA sequences, although quantitative comparisons within narrowly defined sequence windows are feasible (Boswell *et al*, 2017). Thus, direct quantification of the putative N/C gradient of TDP-43 RNA binding sites might require development of new methodological approaches.

### Biophysical regulation of TDP-43—RNA binding

We observed a drastic reduction in TDP-43 passive nuclear exit from permeabilized cell nuclei at 4°C (**Fig 2B-C**), far beyond the approximately two-fold decrease in the passive nuclear import of ERK2-GFP at 4°C in a similar permeabilized cell system (Whitehurst *et al*, 2002), Thus, temperature-dependence of free diffusion alone is unlikely to fully account for the slowing we observed at 4°C. The ability of 1,6-hexanediol to elicit TDP-43 efflux at 4°C (**Fig 2D-E**) suggests that the FG-repeat permeability barrier is temperature-sensitive and likely contributes to the hindrance of TDP-43 passive export at 4°C. Interestingly, TDP-43 nuclear efflux induced by RNA degradation or GU-rich oligomers was also delayed at 4°C, suggesting temperature sensitivity of TDP-43 – RNA binding, which remains to be experimentally verified. RNA-protein interactions generally involve dynamic rearrangements of both binding partners and their stabilization in the complex (Corley *et al*, 2020), suggesting that increased mobility of the complex at a higher temperature may promote TDP-43—RNA dissociation.

Consistent with this explanation, molecular dynamics and structural studies indicate that van der Waals forces support the majority of the TDP-43—RNA interaction (Sun *et al*, 2021) as well as the intramolecular interactions of RRM1—RRM2, which have a significant influence on TDP-43—RNA affinity (Lukavsky *et al*, 2013). Since van der Waals forces steeply decline with distance of the interacting atoms, a temperature-dependent increase in molecular fluctuation thus could stochastically promote TDP-43 unbinding from nuclear RNAs at 37°C.

The repertoire of TDP-43 GU-rich RNA binding partners identified by CLIP-seq is structurally diverse (Polymenidou *et al*, 2011; Tollervey *et al*, 2011), and *in vitro* measurements show that TDP-43 binds to different GU-rich RNAs with widely ranging dissociation constants (~3-3000 nM), with the affinity increasing with the number of perfect GU repeats (Bhardwaj *et al*, 2013). The existence of multiple low affinity-binding sites might have a physiological role in TDP-43 auto-regulation (Avendaño-Vázquez *et al*, 2012; Bhardwaj *et al*, 2013) and helps to explain the ability of short GU-rich oligomers, which bind TDP-43 *in vitro* with low-nM affinity (Bhardwaj *et al*, 2013; French *et al*, 2019), to displace TDP-43 from endogenous RNAs (**Fig 4D,I**). Although RNase caused a near-complete evacuation of TDP-43 from nuclei in permeabilized cells (**Fig 4B-C**), TDP-43 efflux induced by transcriptional blockade (**Fig 1C, Suppl Fig 1C,H,J**) and GU-rich ‘decoy’ oligomers (**Fig 4I**) showed a time- and dose-dependent plateau in living cells. In addition to ongoing active nuclear import, the existence of an ‘export-resistant’ nuclear pool of TDP-43 may result from its binding to other nuclear RNAs such as long non-coding RNAs, which constitute a subset of TDP-43 RNA binding partners (Polymenidou *et al*, 2011; Tollervey *et al*, 2011) and have been shown to play a role in TDP-43 nuclear LLPS (NEAT 1) (Wang *et al*, 2020) and nuclear localization (Malat1) (Nguyen *et al*, 2019). ‘Export-resistant’ TDP-43 may also represent TDP-43 bound to chromatin or nuclear matrix constituents, as supported by TDP-43 chromatin fractionation data (Ayala *et al*, 2008). Since TDP-43 preferentially binds ssDNA (Buratti & Baralle, 2001), direct binding to genomic DNA is uncertain, but TDP-43 could be indirectly associated with chromatin via protein-protein interactions with histones or other chromatin-associated proteins (Freibaum *et al*, 2010). Indeed, TDP-43 was identified among numerous chromatin-associated RBPs in a recent crosslinking/co-precipitation analysis (Rafiee *et al*, 2021). Nevertheless, a large fraction of endogenous, wild-type TDP-43 in our model systems exhibited nuclear RNA concentration-dependent shuttling.

### Other physiologic and pathologic factors that may regulate TDP-43 shuttling

Together with NLS-mediated active nuclear import, our results indicate that TDP-43 RRM1,2-dependent pre-mRNA-binding plays a critical role in establishing the steep concentration gradient of TDP-43 across the nuclear envelope (**Fig 6**). Multiple additional physiologic and pathologic processes likely function in parallel to modulate TDP-43 steady-state localization and availability for active or passive nucleocytoplasmic transport. These include oligomerization (Mompeán *et al*, 2017; Afroz *et al*, 2017; French *et al*, 2019), LLPS (Molliex *et al*, 2015; Conicella *et al*, 2016; Zacco *et al*, 2018; Mann *et al*, 2019; Conicella *et al*, 2020; Carter *et al*, 2021), alternative splicing to truncated isoforms (Weskamp *et al*, 2020), post-translational modifications, such as phosphorylation, ubiquitination, SUMOylation, acetylation, and C-terminal cleavage (Prasad *et al*, 2019; François-Moutal *et al*, 2019), and pathologic aggregation in disease (Arai *et al*, 2006; Neumann *et al*, 2006). Aberrant maturation of TDP-43 LLPS condensates by seeding (Gasset-Rosa *et al*, 2019) or optogenetic clustering (Mann *et al*, 2019) has been shown to progressively divert the soluble TDP-43 pool into insoluble, high molecular weight cytoplasmic aggregates that seemingly become irreversibly unavailable for nuclear transport. Interestingly, GU-RNA oligomers attenuate recombinant TDP-43 aggregation *in vitro* (French *et al*, 2019) and optogenetically-induced cytoplasmic TDP-43 aggregation in living cells (Mann *et al*, 2019), consistent with the notion that access to GU-rich RNAs promotes TDP-43 availability for nucleocytoplasmic shuttling.

### Disruption of RNA metabolism in ALS/FTD

The observation that GU-rich nuclear pre-mRNAs critically regulate TDP-43 nuclear localization raises the possibility that disruption of nuclear RNA metabolism could contribute to TDP-43 nuclear clearance in ALS/FTD. Indeed, disruption of RNA processing and RNA-based therapy development are of growing interest in the neurodegenerative disease field (Nussbacher *et al*, 2019; Butti & Patten, 2019; Zaepfel & Rothstein, 2021). The status of the GU-rich RNA gradient in TDP-43-mislocalized cells in ALS/FTD is unknown, and few studies to date have examined factors predicted to alter the availability of nuclear GU-rich pre-mRNA binding sites, such as transcription and splicing dynamics or nuclear RNA turnover. However, analysis of genome-wide RNA stability in fibroblasts and induced pluripotent stem cells using metabolic labeling (Bru-seq) demonstrated RNA destabilization in ALS patient-derived cells versus controls (Tank *et al*, 2018). Spliceosomal machinery has also appeared as a common denominator in both knockout and interactome analyses of RBPs implicated in familial ALS/FTD (Chi *et al*, 2018a, 2018b). Further investigation of these factors is warranted as well as N/C transcriptome compartmentalization, which has shown changes in nuclear RNA retention in the context of neuropsychiatric disoders (Price *et al*, 2019), and may provide clues as to perturbation of RNA processing in ALS/FTD. Based on the similar behavior of TDP-43 in HeLa cells and neurons following transcriptional inhibition (**Fig 1 and Suppl fig 1**), it appears likely that the RNA-based regulation of TDP-43 nuclear localization is conserved across somatic and neuronal cells. Moreover, it is tempting to speculate that neuron-specific perturbations in RNA processing/metabolism (Mauger *et al*, 2016; Hermey *et al*, 2017; Jaffrey & Wilkinson, 2018; Furlanis *et al*, 2019; Saini *et al*., 2019; Tyssowski & Gray, 2019; Ling *et al*, 2020) may contribute to cell-specific TDP-43 mislocalization in disease.

## Conclusion

In summary, nuclear GU-RNA binding and abundance regulate TDP-43 steady-state nuclear localization by dictating availability for passive nuclear export. Further investigation is needed to determine if disruption of RNA metabolism or localization contribute to TDP-43 nuclear clearance in ALS/FTD. Our findings also suggest that RNA-based approaches may be a useful strategy to restore TDP-43 nuclear localization and attenuate TDP-43 nuclear loss of function defects in disease.

## Methods

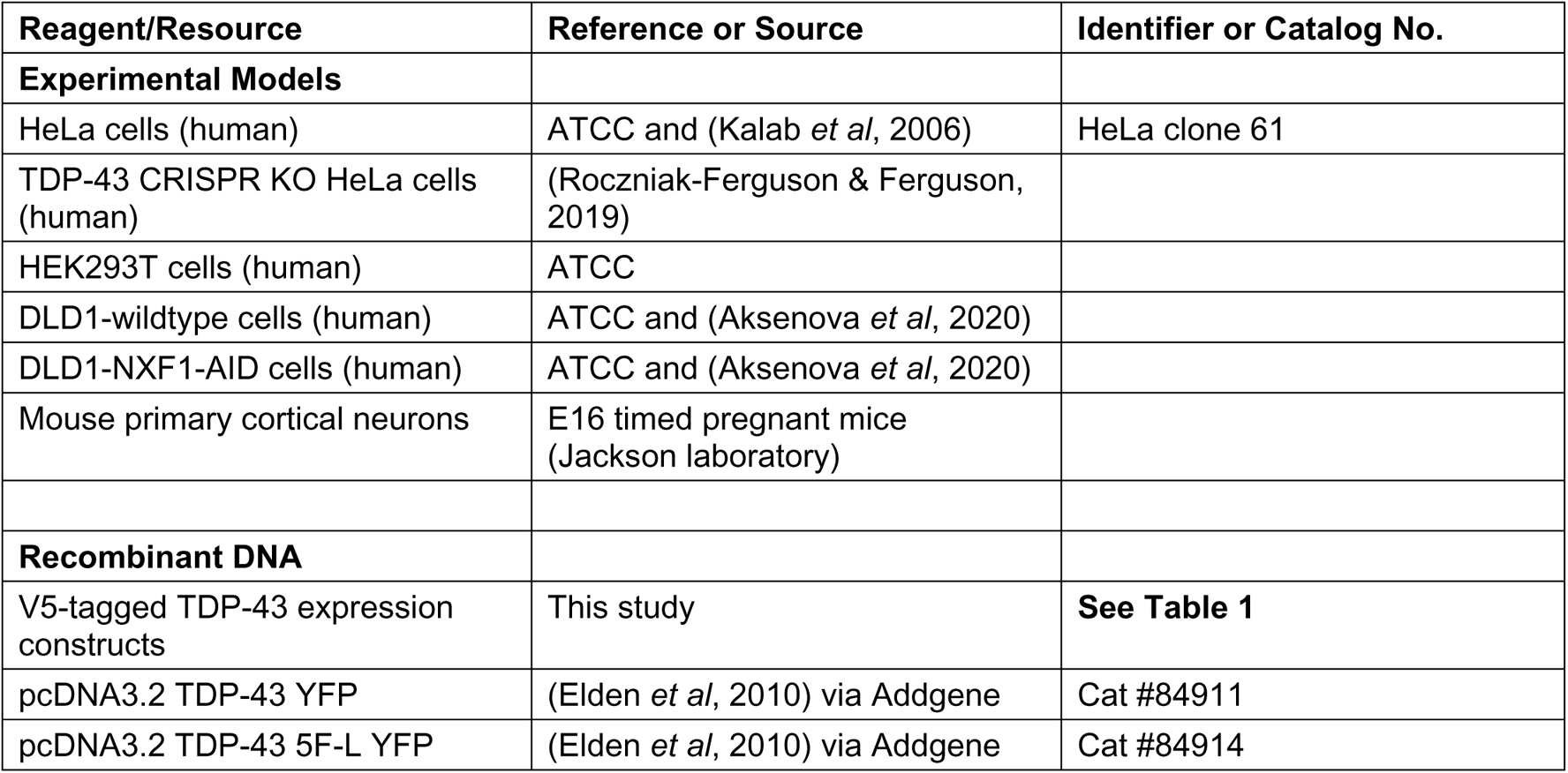
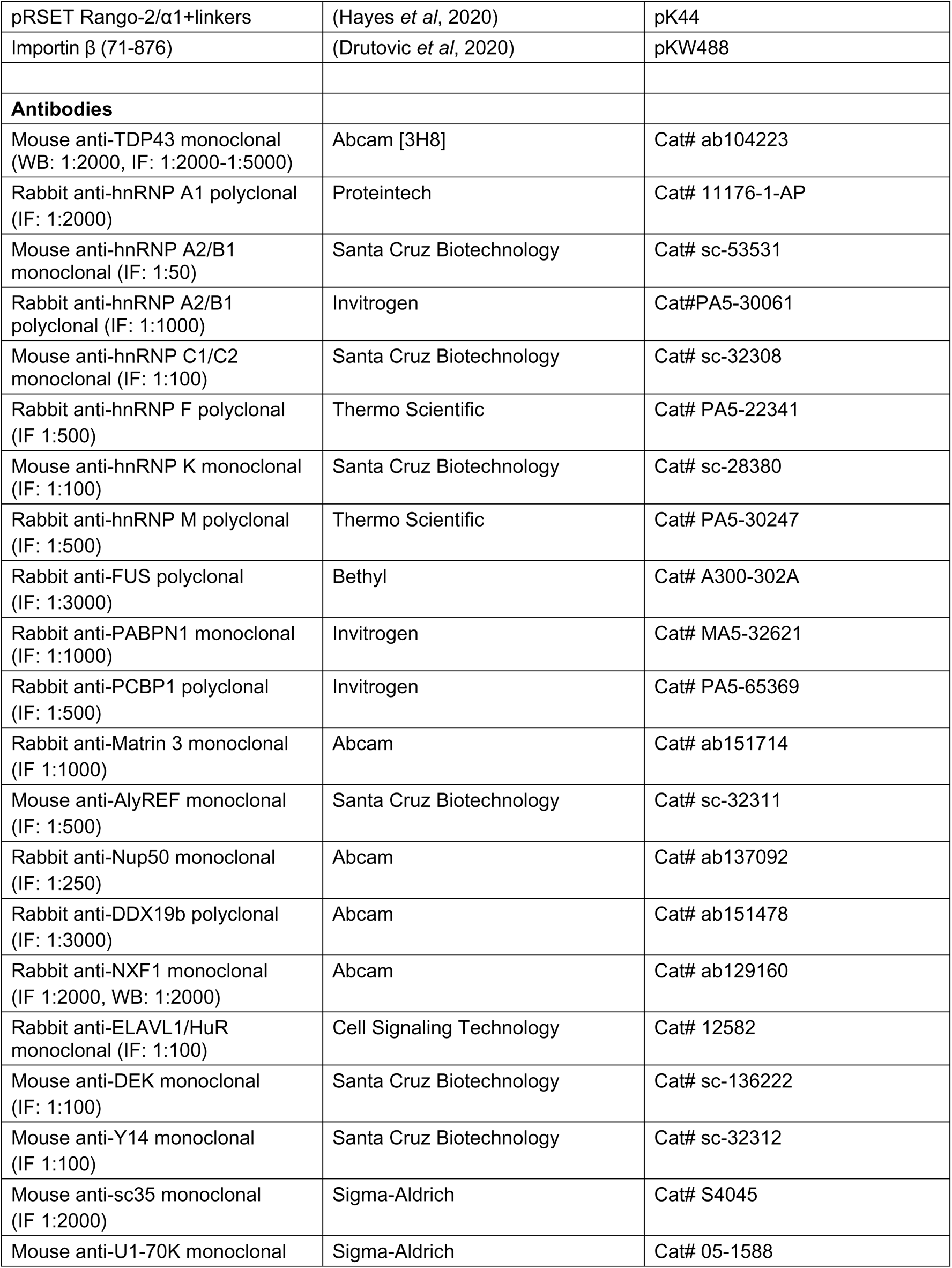
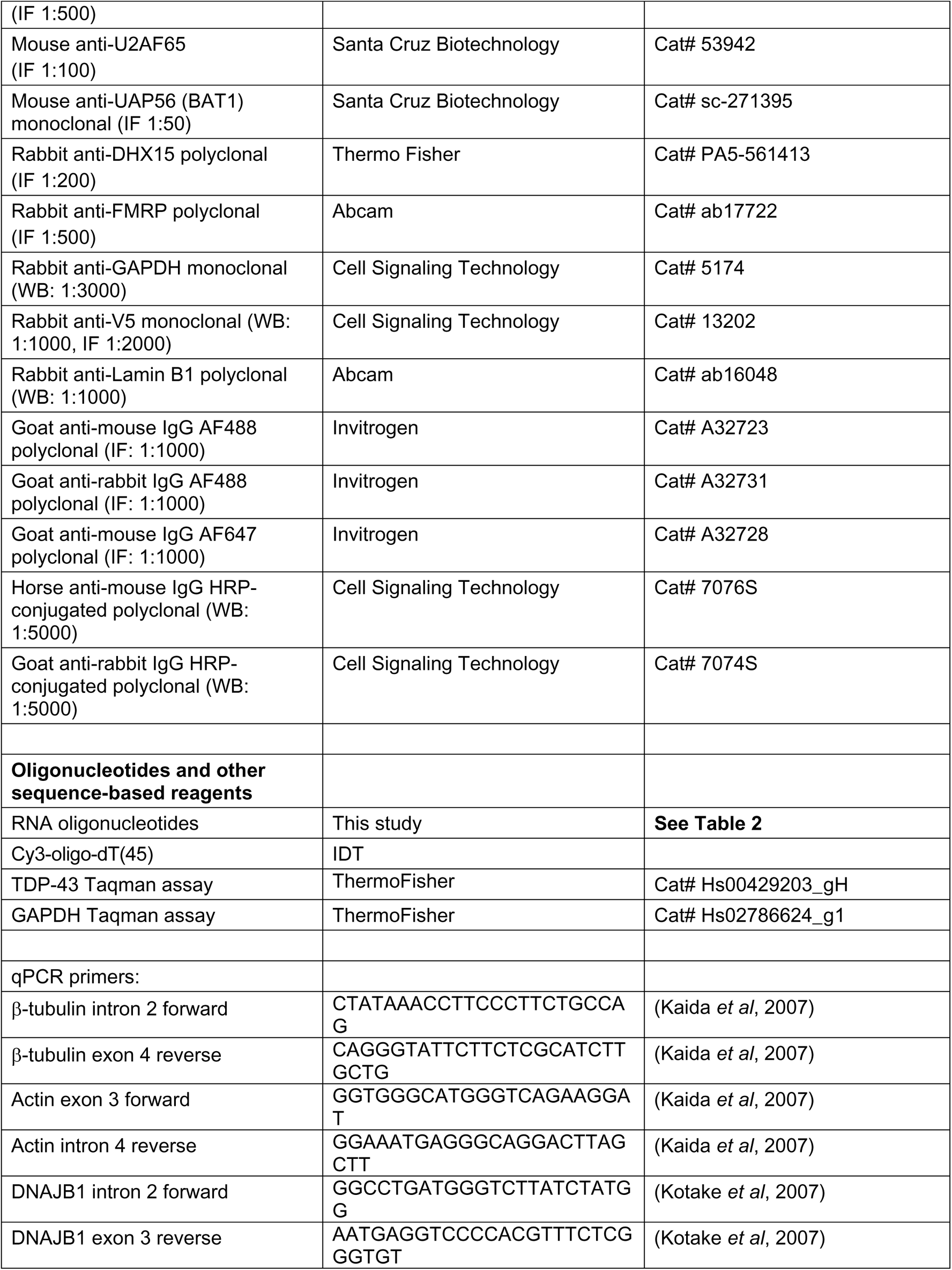
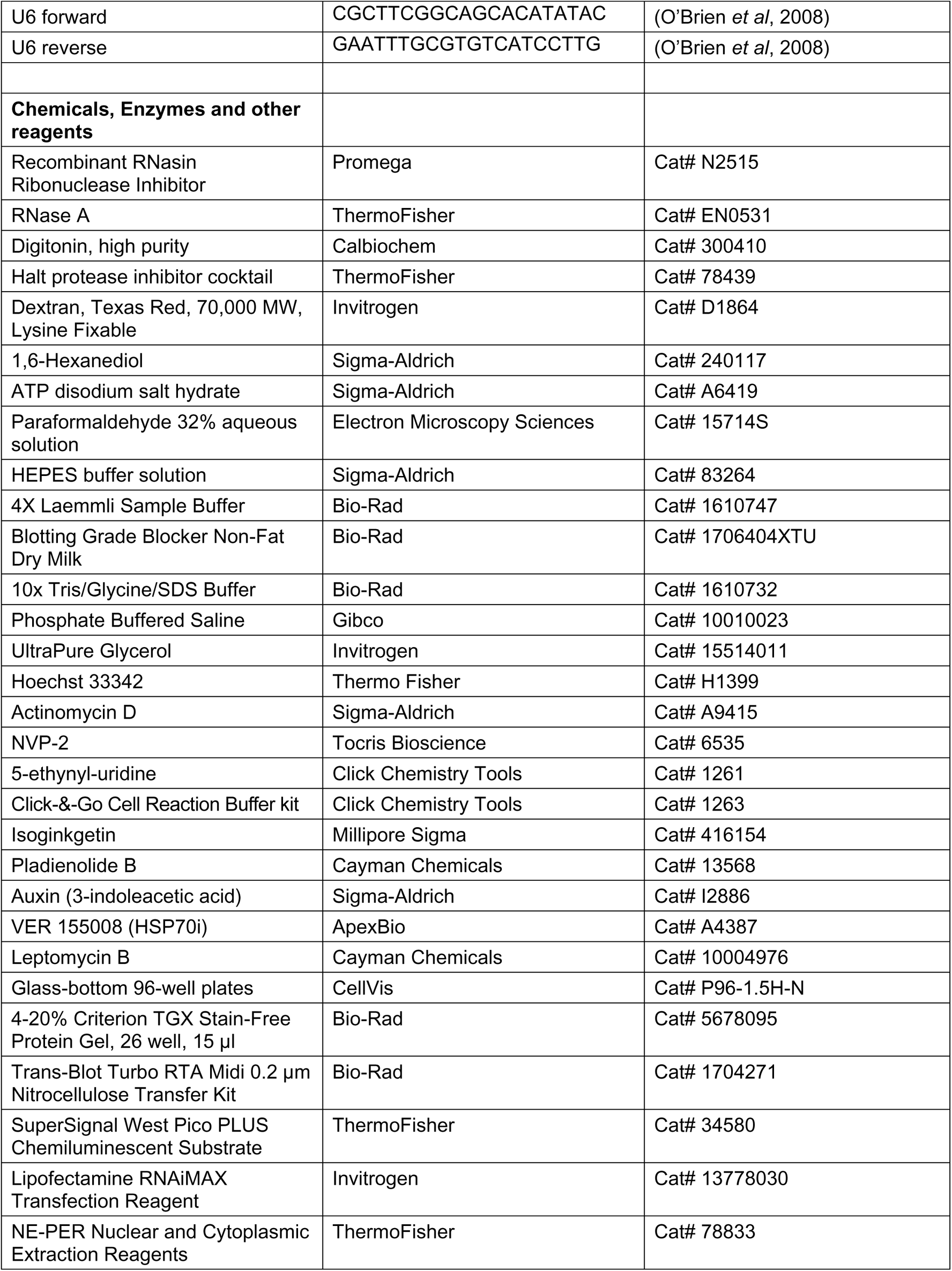
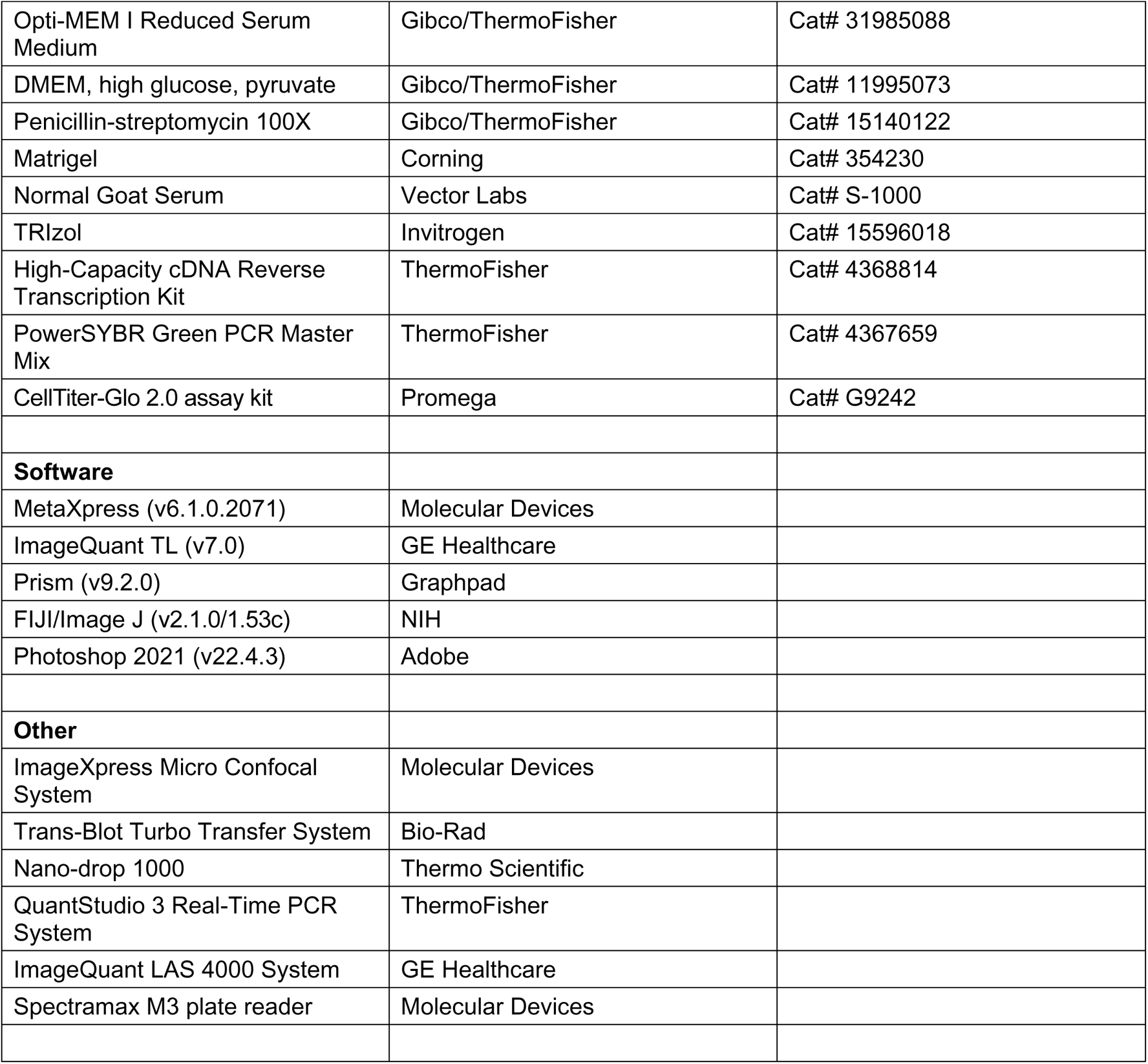

### Immortalized cell culture

A single cell-derived clone of HeLa cells (ATCC) was maintained in OptiMEM (Gibco/ThermoFisher) with 4% FBS and penicillin-streptomycin. HEK293T cells (ATCC) and a monoclonal TDP-43 CRISPR-depleted HeLa cell line (a generous gift from Shawn Ferguson (Roczniak-Ferguson & Ferguson, 2019)), were maintained in DMEM (Gibco/ThermoFisher) with 10% FBS and penicillin-streptomycin. DLD1-wildtype cells (ATCC) and DLD1-NXF1-AID cells (Aksenova *et al*., 2020) were maintained in DMEM (Gibco/ThermoFisher) with 10% FBS and penicillin-streptomycin. All cell lines were validated by STR profiling, routinely verified to be mycoplasma negative, and frequently refreshed from frozen stocks.

### Mouse primary cortical neuron culture

All animal procedures were approved by the Johns Hopkins Animal Care and Use Committee. Timed pregnant C57BL/6J females (Jackson Laboratory) were sacrificed by cervical dislocation at E16, cortex dissociated, and cells plated at 50,000/well on poly-D-lysine/laminin-coated, optical glass-bottom 96-well plates as described (Hayes *et al*, 2020, 2021). Growth medium consisted of Neurobasal supplemented with B27, Glutamax, and penicillin/streptomycin (Gibco/ThermoFisher).

### Cloning of recombinant constructs

Plasmids for the expression of wild-type (WT) human TDP-43 and its variants in tissue culture cells (**Table 1**) were prepared by Twist Biosciences via gene synthesis between the HindIII and NheI sites in the pTwistEF1α expression vector. The expected sequences of TDP-43 open reading frames were verified by Sanger sequencing.

**Table 1.**
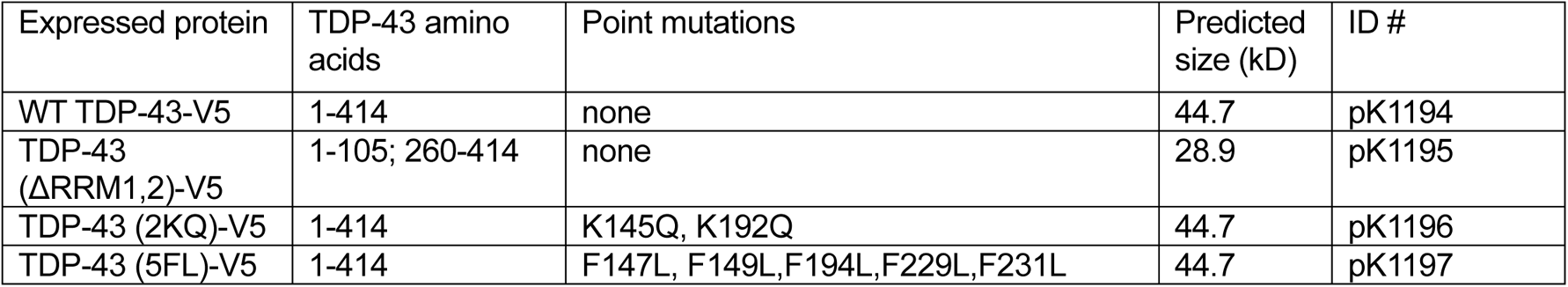
TDP-43 constructs.

| Expressed protein | TDP-43 amino acids | Point mutations | Predicted size (kD) | ID # |
| --- | --- | --- | --- | --- |
| WT TDP-43-V5 | 1-414 | none | 44.7 | pK1194 |
| TDP-43 (ΔRRM1,2)-V5 | 1-105; 260-414 | none | 28.9 | pK1195 |
| TDP-43 (2KQ)-V5 | 1-414 | K145Q, K192Q | 44.7 | pK1196 |
| TDP-43 (5FL)-V5 | 1-414 | F147L, F149L,F194L,F229L,F231L | 44.7 | pK1197 |

### RNA oligonucleotides

Synthetic RNA oligonucleotides (**Table 2**) were obtained from IDT in the form of desalted lyophilized powder. Upon reconstitution in sterile, RNAse and DNAse-free water, single use aliquots were stored at −70°C.

### Permeabilized cell TDP-43 export assays

HeLa cells were plated on Matrigel-coated optical glass-bottom 96 well plates (CellVis), at 12,000-15,000 cells per well targeting 80-90% confluence at 24 h. To permeabilize, cells were rinsed for 2 min in ice-cold PBS, and permeabilized on ice for 10 min in 25-35 ug/mL digitonin (Calbiochem) in transport buffer (TRB, 20 mM HEPES, 110 mM KOAc, 2 mM Mg(OAc)_2_, 5 mM NaOAc, 0.5 mM EGTA, 250 mM sucrose, pH 7.4, with freshly added Halt protease inhibitor (1:100, Promega)). The optimal digitonin concentration varied by cell density and passage number and was optimized prior to each assay such that the majority of plasma membranes were permeabilized while maintaining nuclear exclusion of a 70 kD fluorescent dextran (ThermoFisher). The permeabilized cells were washed twice in ice-cold TRB prior to initiating the export assay.

Assay components, including 1,6-hexanediol (Sigma-Aldrich), ATP (pH 7.5, Sigma-Aldrich), RNase A (ThermoFisher), or RNA oligomers (**Table 2**, synthesized by IDT or Sigma-Aldrich) were premixed in TRB in 96-well plates and equilibrated at the appropriate temperature (4°C, 25°C, or 37°C) prior to cell permeabilization.

With the exception of RNase A-containing assays, TRB was routinely supplemented with RNasin Ribonuclease Inhibitor (1000 U/mL, Promega). Following permeabilization, the assay mix was transferred onto permeabilized cells via multichannel pipette and export was allowed to proceed at the temperature (4°C, 25°C, or 37°C) and for the time (30-60 min) indicated in the figure legends, prior to fixation in 4% paraformaldehyde/PBS (Electron Microscopy Sciences) for 15 min. For each replicate, an additional plate of cells was fixed immediately post-permeabilization, designated ‘time 0’, for the purpose of data normalization across independent biological replicates.

### Permeabilized cell Rango nuclear import assay

Synthesis and nuclear import of the recombinant Rango sensor was carried out as recently described (Hayes *et al*, 2020, 2021). Briefly, following HeLa cell permeabilization as above, Rango nuclear import assays were carried out with or without 2.5 mM ATP (pH 7.5, Sigma-Aldrich) and HEK whole cell lysate (2.5 mg/mL in TRB), at 37°C for 30 minutes. A subset of assays included recombinant human importin β (71-876), a RanGTP-resistant variant (Drutovic *et al*, 2020). Cells were fixed for 15 min in 4% paraformaldehyde/PBS, washed 2x with PBS containing Hoechst (1:5000), and transferred to 50% glycerol/PBS for immediate imaging.

### Live-cell TDP-43 shuttling assays

HeLa and DLD1 cells were plated in uncoated, optical glass-bottom 96-well plates (CellVis) to achieve ~75% confluence at the time of shuttling assays. For transcriptional inhibition, TDP-43 shuttling was initiated by treatment with actinomycin D (Sigma-Aldrich, in DMSO) or NVP-2 (Tocris Bioscience, in DMSO) at doses/times indicated in the figure legends. For splicing inhibition, cells were pretreated for 4 h with escalating doses of isoginkgetin (IGK, Millipore-Sigma, in DMSO) or pladienolide B (PLB, Cayman Chemicals, in DMSO), prior to addition of 250 nM NVP-2. For NXF1 ablation experiments, DLD1-wildtype and DLD1-NXF1-AID cells were pretreated for 0-8 hrs with 0.5 mM synthetic auxin (3-indoleacetic acid, Sigma, in ethanol), prior to addition of 250 nM NVP-2. At the conclusion of experiments, cells were fixed in 4% paraformaldehyde/PBS (Electron Microscopy Sciences) for 15 minutes prior to immunostaining.

### Immunofluorescence

Paraformaldehyde-fixed cells were rinsed with PBS and simultaneously permeabilized and blocked with 0.1% Triton-X 100 and 10% normal goat serum (NGS, Vector Labs) in PBS for 30 min at room temperature. Primary antibody in 10% NGS/PBS was added to the cells and incubated for 60-90 min at room temperature or overnight at 4°C. Cells were rinsed 2x with PBS and AF488 or AF647-labeled secondary antibody (ThermoFisher) was added in 10% NGS/PBS and incubated for 1 h at room temperature. Cells were rinsed with PBS containing Hoechst 33342 and transferred to 50% glycerol/PBS for imaging.

### RNA labeling

For poly(A)-fluorescence in situ hybridization (FISH), cells were fixed in 10% formaldehyde (Sigma-Aldrich) in PBS for 20 min, permeabilized in 0.1% PBS-Tween for 10 min, and washed 3 times in 1x PBS and 2 times in 2x SSC buffer (Sigma-Aldrich) for five min each. Cells were prehybridized for 1 h in hybridization buffer at 42°C (Thermo), and then incubated in Cy3-oligo-dT(45) probe (IDT, 100 nM in hybridization buffer) overnight at 42°C. Cells were washed with decreasing concentrations of SSC buffer at 42°C (2x, then 0.5x, then 0.1x in PBS, for 20 min each), with Hoechst 33342 nuclear counterstain included in the final wash.

For click-chemistry labeling of nascent RNA synthesis, cells were treated with 200 µM 5-ethynyl-uridine in culture media for 30 min, fixed with 4% paraformaldehyde/PBS for 15 min, permeabilized and blocked in 2% BSA/0.1%Tx-100 in PBS for 30 min, and labeled with 0.5 µM AF488-Picolyl Azide using the Click-&-Go Cell Reaction Buffer kit according to the manufacturers’ instructions (all reagents from Click Chemistry Tools). Cells were subsequently immunolabeled for TDP-43 as described above.

### High content imaging and data analysis

Automated imaging and analysis were carried out using an ImageXpress Micro Confocal high content microscope with MetaXpress software (Molecular Devices) as previously described (Hayes *et al*, 2020, 2021). Briefly, nine non-overlapping fields per well were imaged at 20x (immortalized cells) or 40x (neurons) magnification in spinning disc confocal mode with 60 µm pinhole, with exposures targeting half-maximal saturation (33,000 / 65,536 relative fluorescent units (RFU) in unbinned, 16-bit images). The background-corrected mean and integrated nuclear and cytoplasmic intensities and the nuclear/cytoplasmic (N/C) ratio were calculated using the translocation-enhanced module. Number and morphometry of nuclear puncta were calculated using the granularity module. The Hoechst counterstain was used to identify the nuclear/cytoplasmic boundary and nuclear and cytoplasmic compartments were set several pixels in and outside of the nuclear boundary to avoid edge effects. All image analysis was carried out on raw, unaltered images. Raw data were uniformly filtered to exclude errors of cell identification (probe intensity = 0) and non-physiologic N/C ratios (N/C <0.1 or >100). For transient transfections, data were filtered by mean intensity to exclude untransfected cells (<1000 RFU) and highly expressing cells (>45,000 RFU). Resulting data were normalized across technical and biological replicates as percent untreated controls or percent time 0, as indicated in the figure legends. The mean number of cells/well across replicates is also provided. The number of biological replicates (independent cell passages/experiments) was used as the N for statistical analyses.

### qPCR

Cells were rinsed in PBS and lysed in TRIzol (ThermoFisher). Total RNA was isolated following the manufacturer’s protocol and resuspended in nuclease-free water. First-strand cDNA synthesis was performed with 1ug of total RNA using the High-Capacity cDNA Reverse Transcription Kit (ThermoFisher). qPCR was performed using PowerSYBR Green PCR Master Mix (ThermoFisher) with 10ng of cDNA per well and exon-intron primers (500nM) as listed below, using a QuantStudio 3 Real-Time PCR System.

### Nuclear/cytoplasmic fractionation and immunoblotting

24-48 h post-transfection with V5-tagged TDP-43 and RRM mutants, TDP-43 CRISPR KO HeLa cells were lysed for N/C fractionation and SDS-PAGE with the NE-PER kit (ThermoFisher) according to the manufacturer’s instructions. Total protein concentration was measured using the DC Protein Assay kit (Bio-Rad). Nuclear (5 µg total protein) and cytoplasmic (10 µg total protein) fractions were boiled in Laemmli buffer (Bio-Rad), run on Criterion 4-20% Gels (Bio-Rad), and transferred to nitrocellulose membranes using a TransBlot Turbo system (BioRad). Membranes were blocked with 5% non-fat milk in TBS-Tween and probed by sequential incubation in primary antibody overnight at 4°C. Detection was via HRP-conjugated secondary antibodies/chemiluminescence using an ImageQuant LAS 4000 system (GE). Band intensities were measured using ImageQuant software.

### ATP quantification

Permeabilized and live cells in TRB were lysed and ATP levels quantified using the CellTiter-Glo luminescence assay (Promega) according to the manufacturers’ instructions. Luminescence was measured using a SpectraMax M3 microplate reader (Molecular Devices).

### Image processing for figures

Immunofluorescence images were cropped and minimally processed for figures using Adobe Photoshop 2021(v22.4.3) as follows and as indicated in the figure legends. For the purpose of raw intensity comparisons, including the nascent RNA signal (**Fig 1** and **Suppl fig 1**) and the TDP-43 nuclear signal in permeabilized cells (**Fig 2, 4**), the intensity histogram of the designated control was maximized based on the brightest and dimmest pixels, and those parameters were subsequently applied to all other images. For comparisons of shifts in the N/C ratio (all other figures), the intensity histogram was independently maximized according to the brightest and dimmest pixels in each image, to enable adequate visualization of both the nuclear and cytoplasmic compartments. To aid in data visualization, in selected images the fire pseudo-color LUT was applied using FIJI/ImageJ (v2.1.0/1.53c) and the quantitative/linear map is provided in the image. All adjustments were linear (no gamma changes) and applied equally to the entire image. Immunoblots were cropped for space and no other processing was applied. Uncropped immunoblot images are provided in the source data.

### Statistical analysis

Graphing and statistical analyses were carried out using Prism v9.2.0 (Graphpad), according to the methods detailed in the figure legends.

## FIGURE LEGENDS

**Supplemental figure 1.**
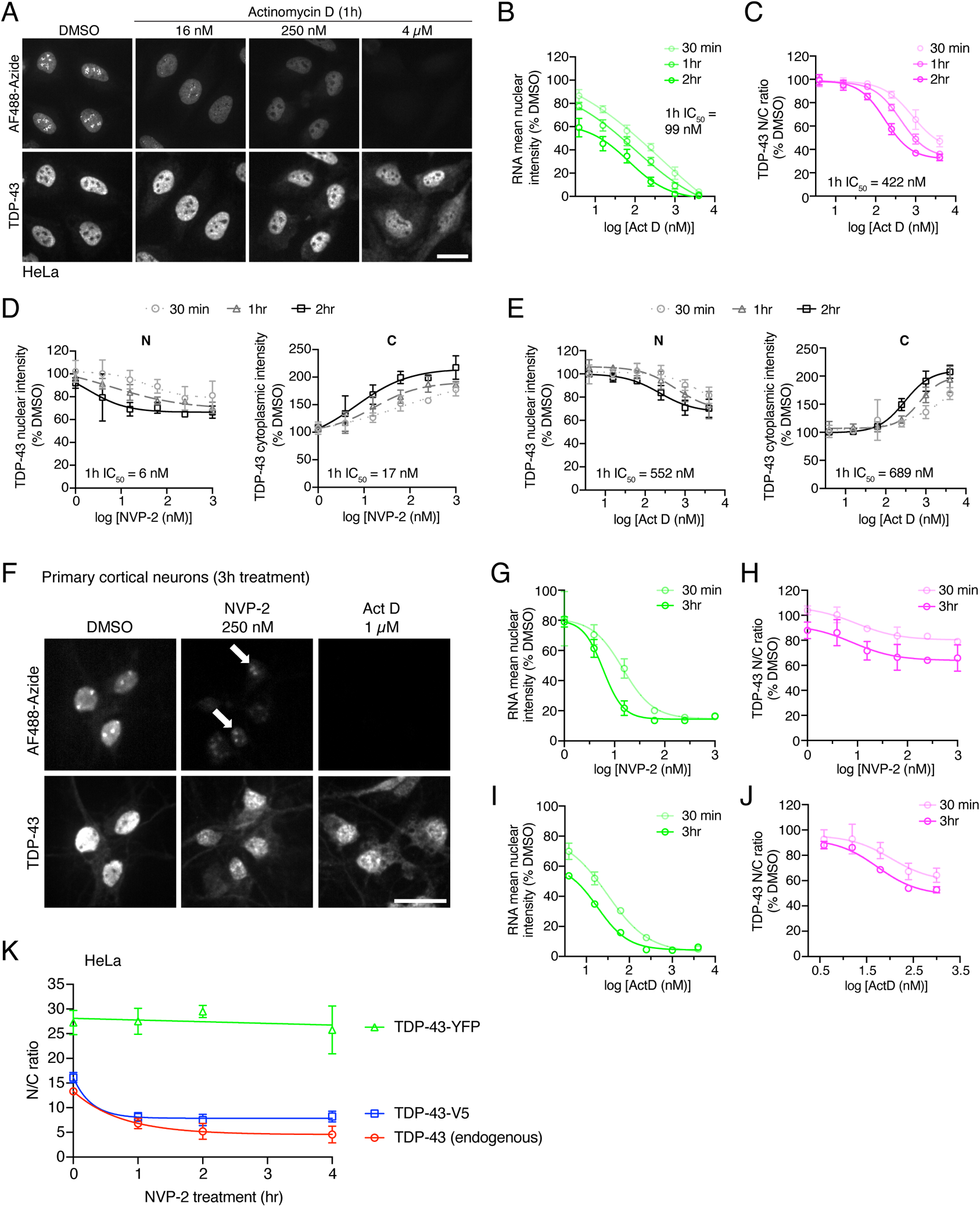
Transcriptional blockade-induced TDP-43 nuclear export in HeLa and neurons. A. Nascent RNAs labeled with AF488-picolyl azide via 5-EU incorporation/‘click chemistry’ (top) and TDP-43 immunofluorescence (bottom) in HeLa cells treated with DMSO (vehicle) vs. Actinomycin D for 1 h. Scale bar = 25 µm. B, C. High content analysis of nascent RNA (AF488-azide) mean nuclear intensity (B) and TDP-43 nuclear to cytoplasmic (N/C) ratio (C) expressed as percent DMSO control. N= mean of 2540 cells/well. Mean ± SD of 3-4 biological replicates per condition is shown. The IC_50_ for 1 h treatment is indicated, as calculated by non-linear regression. D, E. Mean nuclear vs. cytoplasmic intensity of TDP-43 following NVP-2 (D) or Actinomycin D (E) treatment. These are the source data that were used to calculate the N/C ratios in **Fig 1B,C** and **Suppl figure 1B,C**. The IC_50_ for 1 h treatment is indicated, as calculated by non-linear regression. F. Nascent RNAs labeled with AF488-picolyl azide via 5-EU incorporation/‘click chemistry’ (top) and TDP-43 immunofluorescence (bottom) in mouse primary cortical neurons cells treated with DMSO (vehicle), 250 nM NVP-2, or 1 µM Actinomycin D for 3 h. Arrows indicate rRNA puncta unaffected by NVP-2. Scale bar = 25 µm. G-J. High content analysis of nascent RNA (AF488-azide) mean nuclear intensity and TDP-43 nuclear to cytoplasmic (N/C) ratio in primary neurons treated with NVP-2 (G,H) or Actinomycin D (I,J), expressed as percent DMSO control. N= mean of 534 cells/well in each of of 2 biological replicates. Mean ± SD is shown. K. Raw N/C ratio of endogenous TDP-43 vs. TDP-43-YFP and TDP-43-V5 in transiently transfected HeLa cells following 1-4 h treatment with 250 nM NVP-2. N= mean of 2070 cells/well in each of of 3 biological replicates. Mean ± SD is shown.

**Supplemental figure 2.**
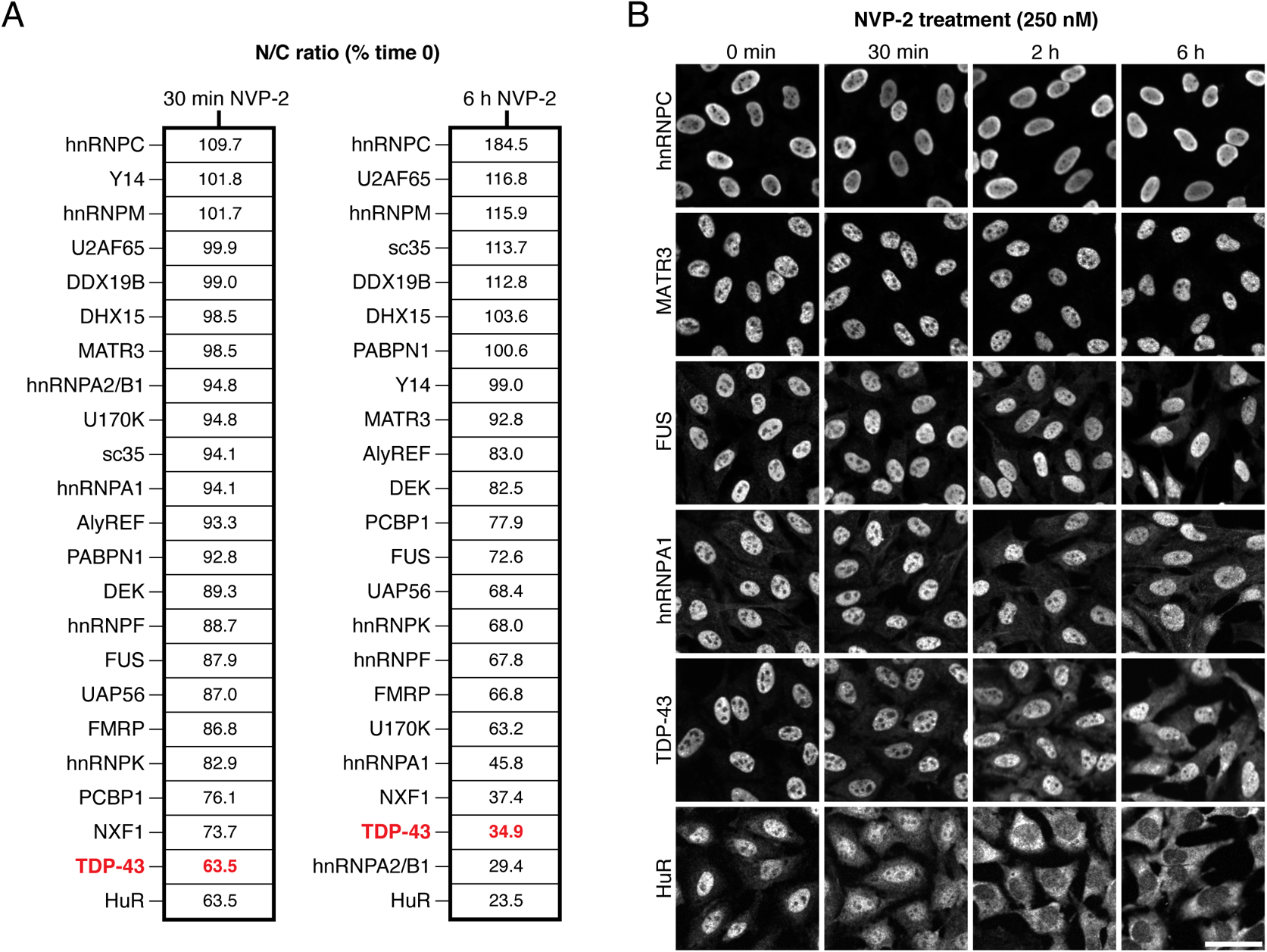
Comparison of NVP-2-induced shuttling among RBPs. A. RBP N/C ratios in HeLa cells after 30 min (left) or 6 h (right) of 250 nM NVP-2 treatment, expressed as % time 0. N= mean of 2716 cells/well. Mean of 2-4 biological replicates per protein is shown. B. RBP immunostaining in NVP-2-treated HeLa cells at designated timepoints following NVP-2 exposure. Scale bar = 50 µm.

**Supplemental figure 3.**
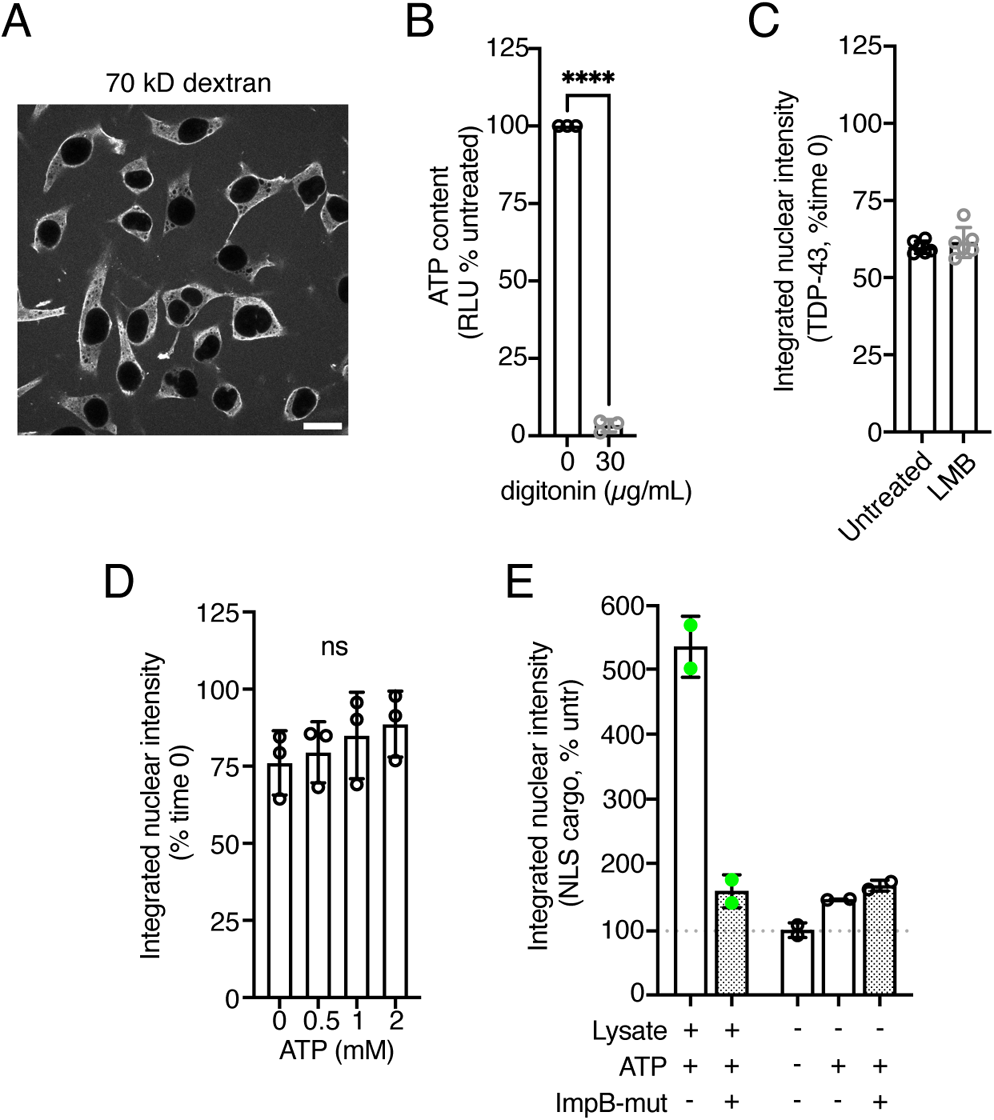
Validation of permeabilized cell passive export assay. A Evaluation of nuclear pore and nuclear membrane integrity by exclusion of 70 kD Texas Red-dextran in digitonin-permeabilized HeLa cells in transport buffer. Scale bar = 20 μm. B. ATP luciferase assay in intact versus digitonin-permeabilized HeLa cells, measured immediately following digitonin permeabilization. N=3 independent biological replicates. ****p<0.0001 by unpaired Student’s t-test. C. TDP-43 integrated nuclear intensity in permeabilized HeLa cells incubated for 30 min at 37°C with or without 100 nM Leptomycin B (LMB) treatment during permeabilization, wash, and export phases. Data are expressed as percent time 0. N = mean of 3005 cells/well in each of 6 technical replicates. D. TDP-43 integrated nuclear intensity in permeabilized HeLa cells incubated for 30 min at 37°C with increasing concentrations of ATP (0 to 2 mM). Data are expressed as percent time 0. N = mean of 2883 cells/well/condition in each of 3 independent biological replicates. NS = not significant, one-way ANOVA with Tukey’s post-hoc test. E. Rango integrated nuclear intensity in permeabilized HeLa cells incubated for 30 min at 37°C with or without HEK cell lysate, ATP (2.5 mM), and dominant-negative importin-β (71-876) mutant. Data are expressed as percent untreated (dotted line). N = mean of 1289 cells/well in each of 2 technical replicates.

**Supplemental figure 4.**
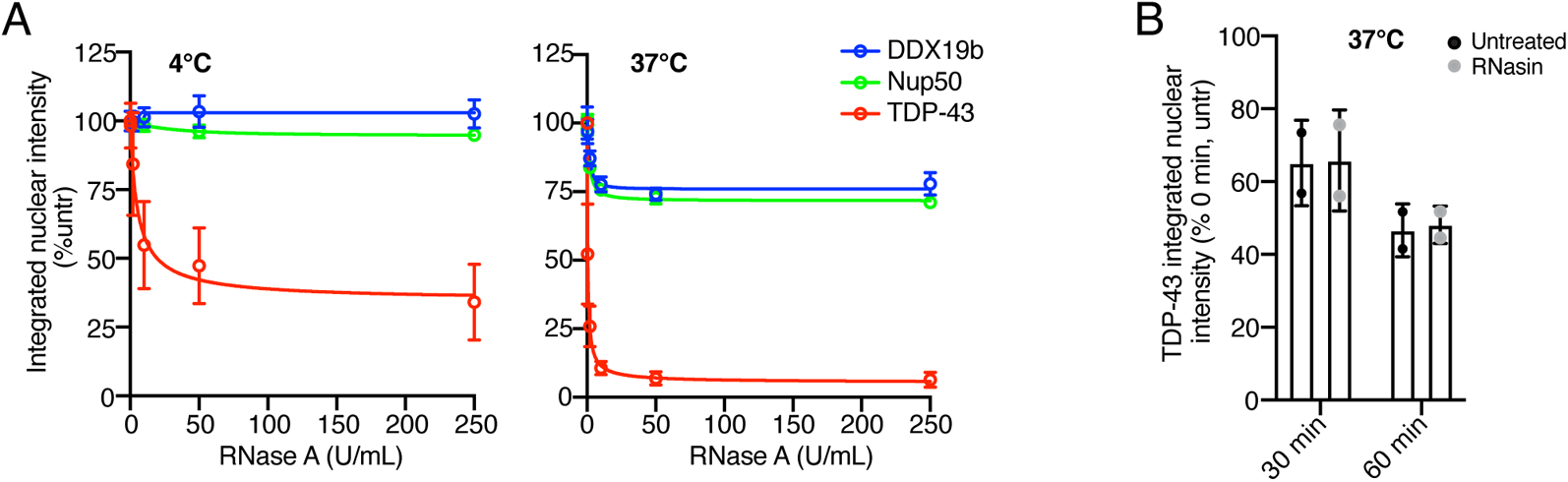
Nuclear and RNA integrity controls. A. Integrated nuclear intensity of DDX19b and Nup50 versus TDP-43 in permeabilized HeLa cells incubated for 30 minutes at 4°C (left) or 37°C (right) with increasing concentrations of RNase A. Data are expressed as percent untreated cells. N = mean of 2868 cells/well in each of 3 technical replicates. Mean ± SD is shown. B. Integrated nuclear intensity of TDP-43 in permeabilized HeLa cells incubated for 30 or 60 min at 37°C with or without RNasin ribonuclease inhibitor. Data are expressed as percent time 0, untreated cells. N = mean of 3262 cells/well in each of 2 independent replicates.

**Supplemental figure 5.**
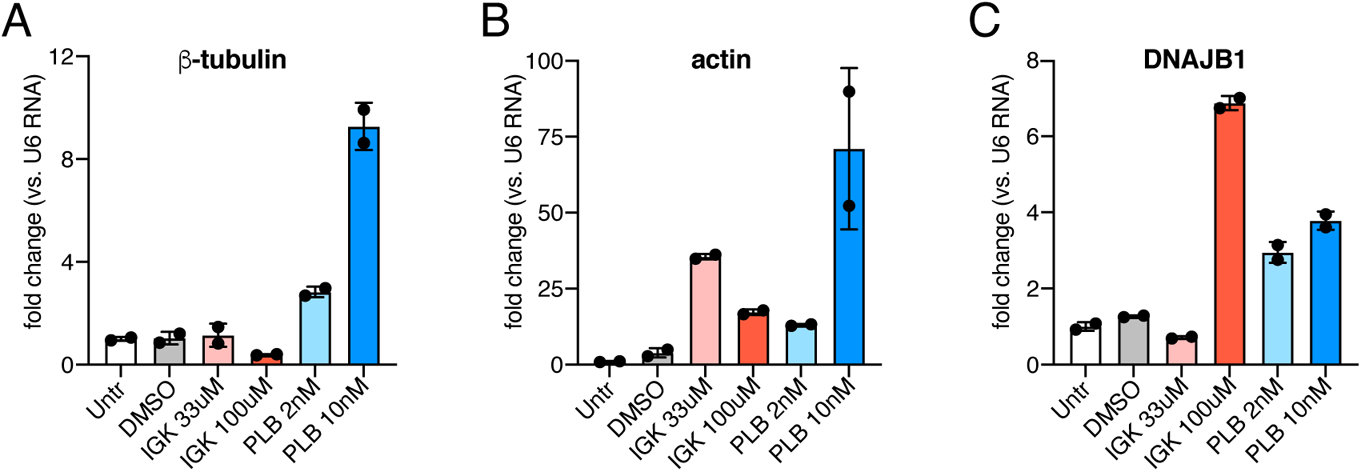
Validation of IGK and PLB-induced intron accumulation. A-C. qRT-PCR quantification of β-tubulin (A), actin (B), and DNAJB1 (C) pre-mRNAs using exon-intron primer pairs, normalized to U6 snRNA expression. HeLa cells were treated for 4 h with IGK or PLB at indicated doses. Mean ± SD is shown for N=2 technical replicates.

**Supplemental figure 6.**
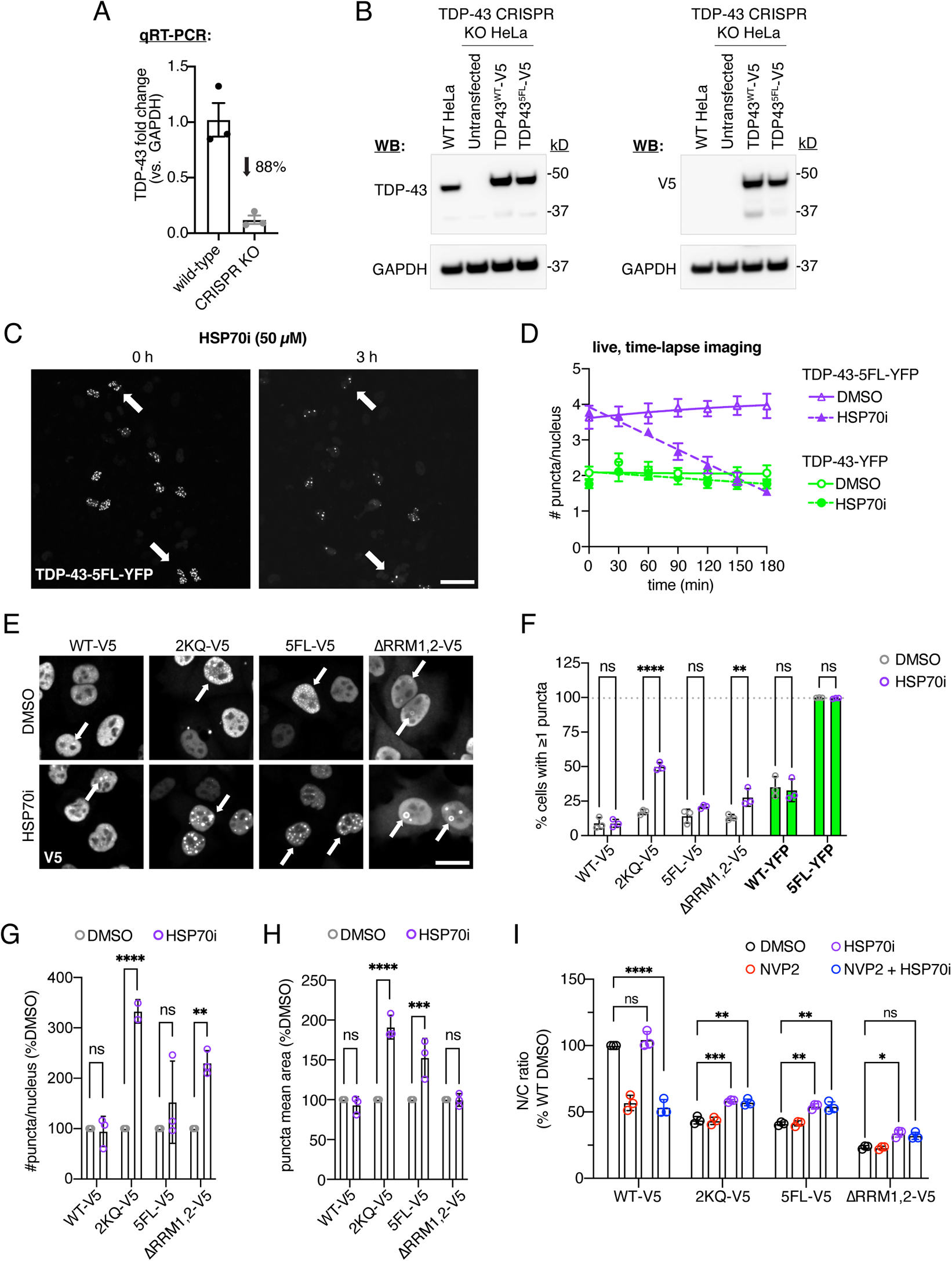
TDP-43 CRISPR KO cell validation and effect of HSP70i on TDP-43 granularity and nuclear localization. A. TDP-43 gene expression in wild-type vs. CRISPR KO HeLa cell lines by qRT-PCR, normalized to GAPDH expression. N=3 technical replicates. B. Immunoblot comparing endogenous TDP-43 protein expression in wild-type vs. CRISPR KO HeLa cells (left). Level of expression of transiently transfected V5-tagged constructs vs. endogenous TDP-43, and immunodetection by TDP-43 (left) and V5 (right) antibodies is also shown. C. YFP epifluorescence in TDP-43 CRISPR KO cells transiently transfected with TDP-43-5FL-YFP, before and after 3 h treatment with HSP70i (50 µM). Arrows highlight examples of reduction in number of nuclear granules. Note that moderate cell migration occurs over the timecourse of the assay. Scale bar = 50 µm. D. High content analysis of number of TDP-43-YFP and TDP-43-5FL-YFP granules per nucleus in DMSO vs. HSP70i-treated cells, assessed via live, time-lapse imaging. N = mean of 2171 cells/well/condition in each of two independent biological replicates. Mean ± SD is shown. E. V5 immunofluorescence in TDP-43 CRISPR KO cells transiently transfected with V5-tagged wild-type and RRM mutant TDP-43 constructs, and treated for 3 h with DMSO vs. HSP70i (50 µM). Note: Rare examples of puncta-containing nuclei are shown (arrows) which constitute only a minority of the cell population as quantified in (F). The intensity histogram for each image was independently maximized across the full range. Given the intense fluorescence of the puncta, this prevents visualization of the N/C gradients which are better appreciated in the more representative images in Fig 6A. Scale bar = 25 µM. F-H. High content granularity analysis in TDP-43 CRISPR KO cells transiently transfected with V5- or YFP-tagged wild-type and RRM mutant TDP-43 constructs, and treated with DMSO or HSP70i (50 µM) for 3h. Parameters include % cells with ≥ 1 puncta (F), # puncta/nucleus (G, expressed as % DMSO), and mean puncta area (H, expressed as % DMSO). N = mean of 1156 cells/well/condition in each of three independent biological replicates. Mean ± SD is shown. I. N/C ratio of V5 immunofluorescence signal, in TDP-43 CRISPR KO cells transiently transfected with V5-tagged wild-type and RRM mutant TDP-43 constructs, and treated with DMSO, NVP2 (250 nM), HSP70i (50 µM), or both for 3h. Data are expressed as percent wild-type DMSO-treated cells. N = mean of 788 cells/well/condition in each of three independent biological replicates. In F-I: ns = not significant, *p<0.05, **p<0.01, ***p<0.001, ****p<0.0001 for designated comparisons by 2-way ANOVA with Sidak’s multiple comparisons test.

## ACKNOWLEDGEMENTS

Lin Xue, Svetlana Vidensky, and Lin Jin provided expert technical assistance. Pei-Hsun Wu provided imaging assistance. Lyudmila Mamedova and Barbara Smith provided administrative support. This research was supported by NIH K08NS104273 and R01NS123538 (L.R.H.), DOD/CDRMP W81XWH1910209 (J.D.R. and L.R.H.), and the Robert Packard Center for ALS Research (L.R.H. and P.K.).

## AUTHOR CONTRIBUTIONS

L.D. – conceptualization, methodology, investigation, validation, formal analysis, visualization, writing-original draft, writing-review & editing; B.Z. – methodology, investigation, writing-review & editing; V.A. – methodology, resources, writing-review & editing; M.D. – resources, supervision, writing-review & editing; J.D.R. – supervision, funding acquisition, writing-review & editing; P.K. – conceptualization, methodology, investigation, resources, visualization, writing-original draft, writing-review & editing, funding acquisition; L.R.H. – conceptualization, methodology, investigation, validation, formal analysis, visualization, writing-original draft, writing-review & editing, supervision, funding acquisition.

## CONFLICTS OF INTEREST

The authors declare that they have no conflicts of interest to report.

